# Programmable Protein Reference Standards for benchmarking sub-10 nm Fluorescence Microscopy

**DOI:** 10.64898/2026.07.27.740971

**Authors:** Made Budiarta, Dominic Helmerich, Marcel Streit, Marvin Jungblut, Sören Doose, Philip Kollmannsberger, Markus Sauer, Gerti Beliu

## Abstract

Fluorescence microscopy is increasingly used to measure molecular organization at length scales where labeling, photophysics and sample preparation can dominate quantitative accuracy. Reference standards are therefore needed that combine defined nanoscale geometry with a protein-like environment and compatibility with biological imaging. Here we introduce circular tandem repeat protein as programmable protein standards for benchmarking sub-10-nm fluorescence microscopy. Using genetic code expansion and bioorthogonal labeling, we generated compact protein rings carrying up to six labeling sites. Photoswitching fingerprint analysis revealed geometry-dependent localization accumulation and blinking kinetics, demonstrating that short-range fluorophore interactions can be assessed as measurable benchmark parameters. DNA-PAINT confirmed accessible docking sites and programmed valency at the single-particle level. We further established recombinant tethering and genetically encoded membrane display, extending the standards to cellular environments. cTRP PicoRulers were also compatible with expansion microscopy. Together, cTRPs provide a modular protein-based platform for evaluating molecular-scale imaging performance in purified and cellular environments.

## 1. Introduction

Super-resolution microscopy (SRM) has transformed cell biology by enabling molecular organization to be visualized beyond the diffraction limit. Recent advances in DNA-PAINT,^1,2^ MINFLUX,^3,4^ Expansion Microscopy (ExM)^5^ and related approaches have pushed fluorescence imaging toward the single-digit nanometer scale.^3,6,7^ As imaging approaches molecular length scales, however, localization precision is no longer the only factor limiting quantitative accuracy. Incomplete labeling can leave subsets of molecules within nanoscale assemblies undetected, while short-range interactions between neighboring fluorophores can alter blinking, brightness and detection probability.^8,9^ In addition, linkage error arising from the physical separation between a fluorophore and its target can become comparable to the biological distances being measured, thereby resulting in false impressions of the underlying true molecular organization.^10,11^ Consequently, molecular structures may be localized with nanometer precision yet still be measured or interpreted inaccurately. These effects are particularly relevant for dense protein assemblies, membrane receptors and nanoscale protein clusters, where differences of only a few nanometers can significantly change biological interpretation. Reference standards with known geometry and controlled fluorophore placement are therefore essential for distinguishing true molecular organization from method-dependent artifacts.

Several classes of reference structures have advanced quantitative super-resolution microscopy. Endogenous cellular structures, such as microtubules and nuclear pore complexes provide biologically relevant benchmarks and have been widely used to assess microscope performance, labeling efficiency and molecular counting.^12,13^ On the contrary, DNA nanotechnology such as DNA origami provide highly programmable geometries and have been central for testing resolution at nanometer length scales.^14^ Although it has been shown that they can be presented or tethered on cell surfaces, DNA origami do not fully reproduce the compact protein environment, genetically encoded labeling chemistry or native membrane anchoring encountered in biological measurements.^15,16^ Protein-based nanorulers could bridge this gap, but existing designs such as PCNA PicoRulers are constrained by fixed symmetry and limited control over fluorophore number and arrangement.^17,18^

We reasoned that circular tandem repeat proteins (cTRPs)^19^ could provide a programmable protein scaffold for quantitative sub-10 nm fluorescence microscopy. These *de novo*-designed proteins form compact ring-shaped architectures of approximately 10 nm diameter and contain repeated surface-exposed loop regions that could be engineered as defined labeling sites (Figs. 1a,d). By combining this architecture with genetic code expansion (GCE) and bioorthogonal click chemistry, fluorophores can be positioned at selected sites within a genetically encoded protein scaffold with minimal linkage error (Figs. 1b,c).

**Figure 1.**
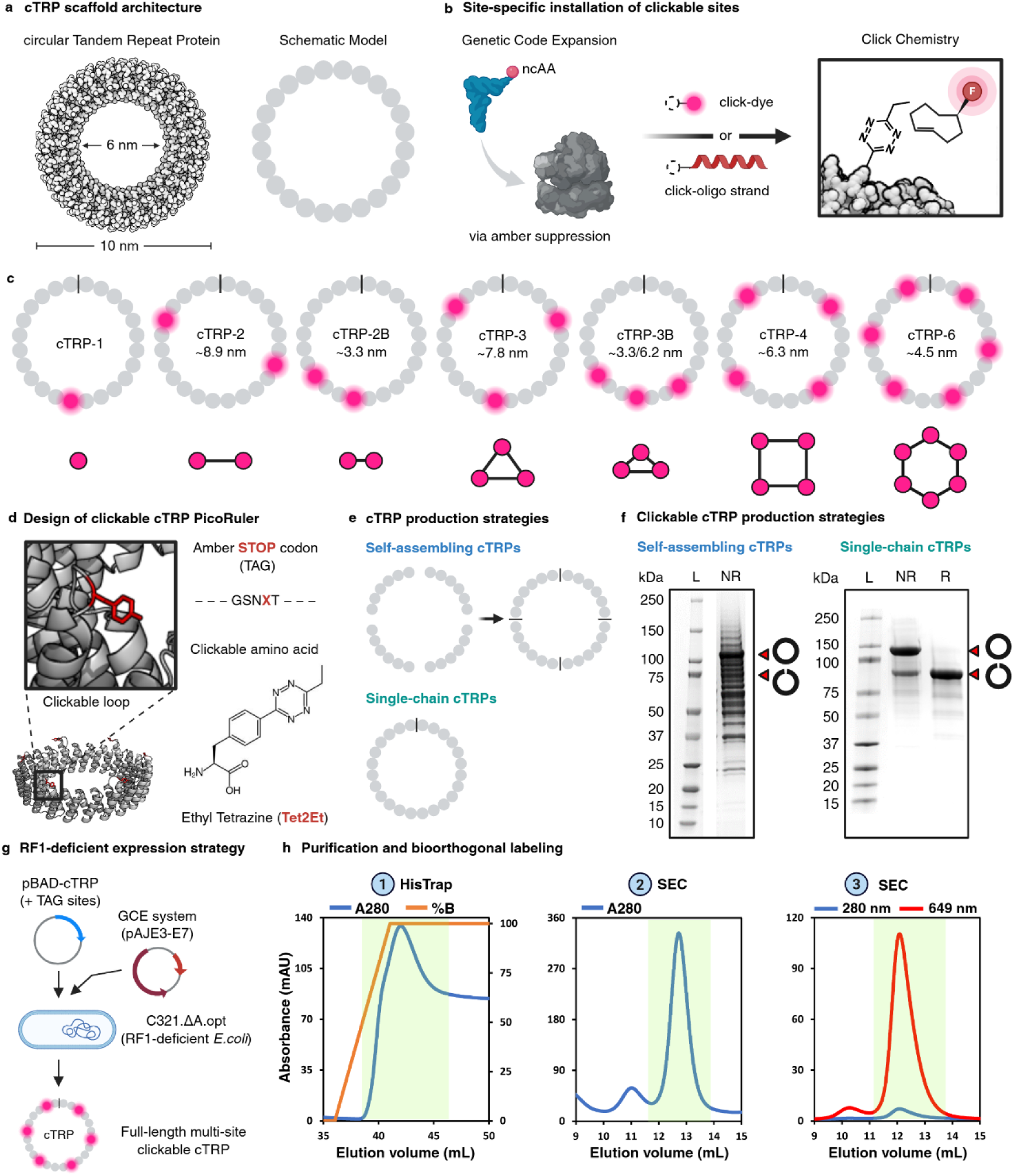
Design and production of programmable cTRP PicoRulers. a, Molecular and schematic representations of the 24-repeat cTRP scaffold. **b,** Site-specific incorporation of a non-canonical amino acid by amber suppression, followed by bioorthogonal labeling with trans-cyclooctene-modified fluorophores or DNA oligonucleotides. **c,** cTRP PicoRulers carrying one, two, three, four or six clickable sites in distributed or compact arrangements. Approximate designed inter-site distances and simplified geometries are shown. **d,** Design of a clickable surface-exposed loop containing Tet2Et within a GSNXT amber-suppression context. **e,** Self-assembling and single-chain cTRP production strategies. **f,** Representative SDS–PAGE analysis of clickable cTRPs. Arrowheads indicate species assigned to closed- and open-ring conformations. L, molecular-mass ladder; NR, non-reducing; R, reducing. **g,** RF1-deficient expression strategy for multisite Tet2Et incorporation. **h,** Representative HisTrap purification, size-exclusion chromatography, Cy5–TCO labeling and subsequent size-exclusion chromatography. Shaded regions indicate pooled fractions.

Here we establish cTRP next generation PicoRulers as programmable protein reference standards for benchmarking sub-10 nm fluorescence microscopy. By combining cTRP design with genetic code expansion and bioorthogonal labeling, we generated ∼ 10-nm protein rings carrying one, two, three, four or six genetically specified labeling sites. The recently developed photoswitching fingerprint analysis revealed geometry-dependent localization accumulation and blinking kinetics as measurable benchmark parameters for assessing short-range fluorophore interactions.^8^ DNA-PAINT provided an orthogonal single-particle readout of docking-site accessibility and programmed valency. We further established recombinant tethering and genetically encoded membrane display, extending the platform to cellular environments, and demonstrated compatibility with expansion-based imaging, including post-expansion labeling and optical readout. Together, cTRP PicoRulers provide a modular protein-based platform for evaluating labeling-site accessibility, fluorophore photophysics and molecular-scale imaging performance in purified and cellular environments.

## 2. Results

### 2.1 Design and production of programmable cTRP reference standards

To generate protein-based reference standards with defined fluorophore number and spacing, we used cTRPs as compact and programmable scaffolds. cTRPs are *de novo*-designed ring-shaped proteins composed of 24 repeated units with an outer diameter of approximately 10 nm (Fig. 1a, Extended Data Fig. 1a),^19^ placing their full geometry within the distance range relevant for sub-10 nm fluorescence microscopy. Each repeat contains surface-exposed loop regions that can be engineered without disrupting the overall architecture (Fig. 1d, Extended Data Fig. 2b), providing a modular framework for positioning fluorophores at selected sites around the ring (Fig. 1c). This architecture allowed us to vary two parameters that are difficult to control independently in biological samples: the number of fluorophores on a molecular object and their relative nanoscale spacing.

We first established recombinant production of wild-type cTRP scaffolds in *E. coli* C41 (DE3). In principle, cTRP rings can be produced either as a single polypeptide chain or through self-assembly of smaller subunits stabilized by disulfide bonds (Fig. 1e, Extended Data Fig. 1a, Supplementary Table S1).^19^ We tested both formats to determine which architecture would best support subsequent multi-site functionalization. Both formats yielded the expected ring architecture, but the self-assembling format required additional purification steps to separate complete rings from partially assembled species (Extended Data Figs. 1b,c). Transmission electron microscopy and SDS-PAGE analysis confirmed formation of the expected ring architecture in both formats (Extended Data Figs. 1d,e).

We next converted cTRPs into clickable reference standards by introducing amber stop codons into selected loop regions and incorporating the non-canonical amino acid (ncAA) ethyl-tetrazine (Tet2Et) via GCE (Figs. 1b,d).^20^ To enable efficient amber suppression, we inserted Tet2Et within a GSNXT motif (X = Tet2Et), in which GS provides a flexible linker and NXT (AAT TAG ACT) supplies a favorable codon context for ncAA incorporation (Fig. 1d, Extended Data Fig. 2b).^21^ We initially used self-assembling cTRPs because this format allowed click-sites valency to be increased through assembly of subunits. Using this strategy, we designed cTRP variants carrying one, two, three, four or six clickable sites around the ring (Extended Data Fig. 2a).

Although self-assembling cTRPs confirmed the feasibility of introducing clickable sites into the scaffold, purification became increasingly inefficient as the number of amber codons increased. Truncated and incompletely assembled products accumulated and interfered with isolation of homogeneous full-ring structures, particularly for higher-valency designs (Fig. 1f, Extended Data Fig. 2a). We therefore shifted to single-chain cTRPs expressed in the RF1-deficient *E. coli* strain C321.ΔA.opt,^22^ which reduces translational termination at amber codons and supports multi-site ncAA incorporation (Fig. 1g). This strategy enabled production of full-length cTRP variants carrying one, two, three, four or six Tet2Et residues at defined positions (Figs. 1c,f,h, Extended Data Fig. 2e, Supplementary Table S2).

The resulting cTRP scaffolds were site-specifically labeled with Cy5-TCO through bioorthogonal click chemistry (IEDDA ligation) and purified by size-exclusion chromatography (Fig. 1h, Extended Data Fig. 2c). Spectroscopic analysis confirmed fluorophore incorporation across the panel, with labeling values consistent with the expected increase in fluorophore valency from cTRP-1 to cTRP-6 (Extended Data Fig. 2d). Collectively, these findings establish that single-chain expression in RF1-deficient bacteria overcomes the main production bottleneck of the self-assembling format and provides a robust route to programmable cTRP reference standards with controlled fluorophore valency and designed nanoscale geometry.

### 2.2 Membrane display extends cTRP PicoRulers to cellular imaging environments

Reference standards for molecular-scale fluorescence microscopy are most informative when they capture the same sources of measurement bias encountered in biological samples. We therefore asked whether cTRP PicoRulers could be tethered at the cell surface, where membrane orientation, local crowding, labeling accessibility and fluorophore interactions can influence quantitative fluorescence readouts in ways that are not reproduced by purified particles immobilized on glass surface or DNA nanotechnology.

We first established a recombinant membrane-tethering strategy. Purified Cy5-labeled cTRP PicoRulers were functionalized with HaloTag ligand through NHS-bioconjugation and then incubated with cells expressing extracellular HaloTag at the plasma membrane (Fig. 2a, Extended Data Fig. 3a). Mobility-shift assays with purified HaloTag protein confirmed successful functionalization as the cTRP PicoRulers formed higher-molecular-weight species after incubation with HaloTag protein, consistent with binding of one or more HaloTag molecules to ligand-modified cTRPs (Fig. 2b). When added to HaloTag-expressing cells, the functionalized cTRPs produced membrane-localized fluorescence, demonstrating that recombinant cTRP PicoRulers can be tethered to the cellular surface (Extended Data Fig. 3b). Colocalization of anti-c-Myc-AF488-labeled extracellular HaloTag with Cy5-labeled cTRP PicoRulers further supported HaloTag-dependent recruitment of recombinant cTRPs to the cell surface.

**Figure 2.**
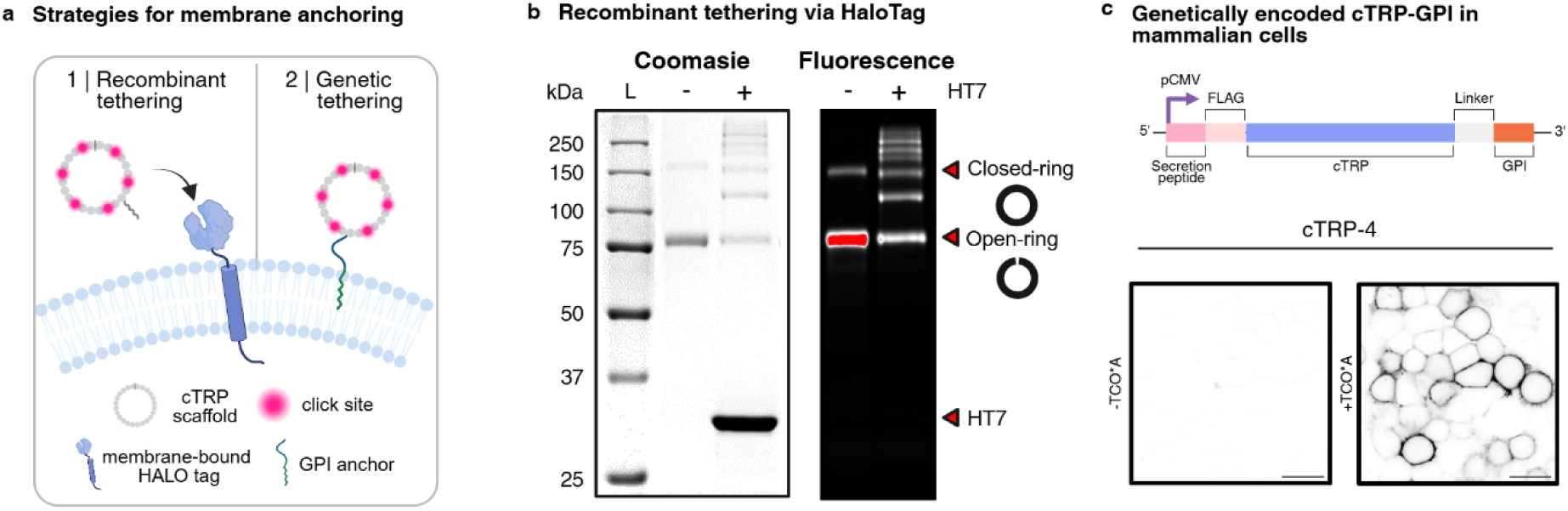
Recombinant and genetically encoded membrane display of cTRP PicoRulers. a, Complementary strategies for cell-surface presentation: purified cTRPs tethered to extracellular HaloTag through a covalently attached ligand, or genetically encoded cTRPs carrying a C-terminal GPI anchor. **b,** Recombinant HaloTag-mediated tethering. Representative SDS–PAGE mobility-shift analysis of Cy5-and HaloTag ligand-functionalized cTRP-2 following incubation with or without purified HaloTag protein (HT7), confirming successful dual functionalization of the cTRP PicoRuler. The gel was imaged under coomassie staining (left) and Cy5 fluorescence (right). **c,** Genetically encoded membrane display. Top, mammalian expression construct comprising a secretion peptide, FLAG epitope, cTRP sequence, linker and GPI-anchor signal. Bottom, representative fluorescence images of cells expressing cTRP-4 in the absence or presence of TCO*A followed by bioorthogonal fluorophore labeling. Scale bars, 20 µm.

We next developed a genetically encoded membrane-display format. cTRP constructs were fused to an N-terminal secretion signal and a C-terminal GPI-anchor sequence, enabling trafficking through the secretory pathway and presentation at the outer leaflet of the plasma membrane (Figs. 2a,c, Supplementary Table S3). Because the bacterial GSNXT amber-suppression motif contains a potential N-linked glycosylation site, we replaced it with a GSGXL motif (GXL = GGG TAG CTC), an optimized codon context for amber suppression in mammalian cells (Extended Data Fig. 4a).^23^ Co-expression with an amber-suppression system in the presence of a clickable ncAA followed by tetrazine-dye labeling produced clear membrane-localized fluorescence signals for cTRP variants carrying different numbers of labeling sites (Fig. 2c, Extended Data Fig. 4b).

### 2.3 Defined cTRP geometries generate distinct photoswitching fingerprints

We next used cTRP PicoRulers to test whether defined nanoscale protein geometries generate distinguishable photoswitching signatures (Fig. 3a). Fluorophores separated by only a few nanometers can interact through energy transfer, leading to quenching and accelerated blinking kinetics, thereby changing localization probability and the temporal accumulation of detected events in single-molecule localization microscopy.^8^ We therefore reasoned that genetically encoded cTRPs with controlled fluorophore number and spacing could serve as a protein-based benchmark for photoswitching fingerprint analysis (PFA).

**Figure 3.**
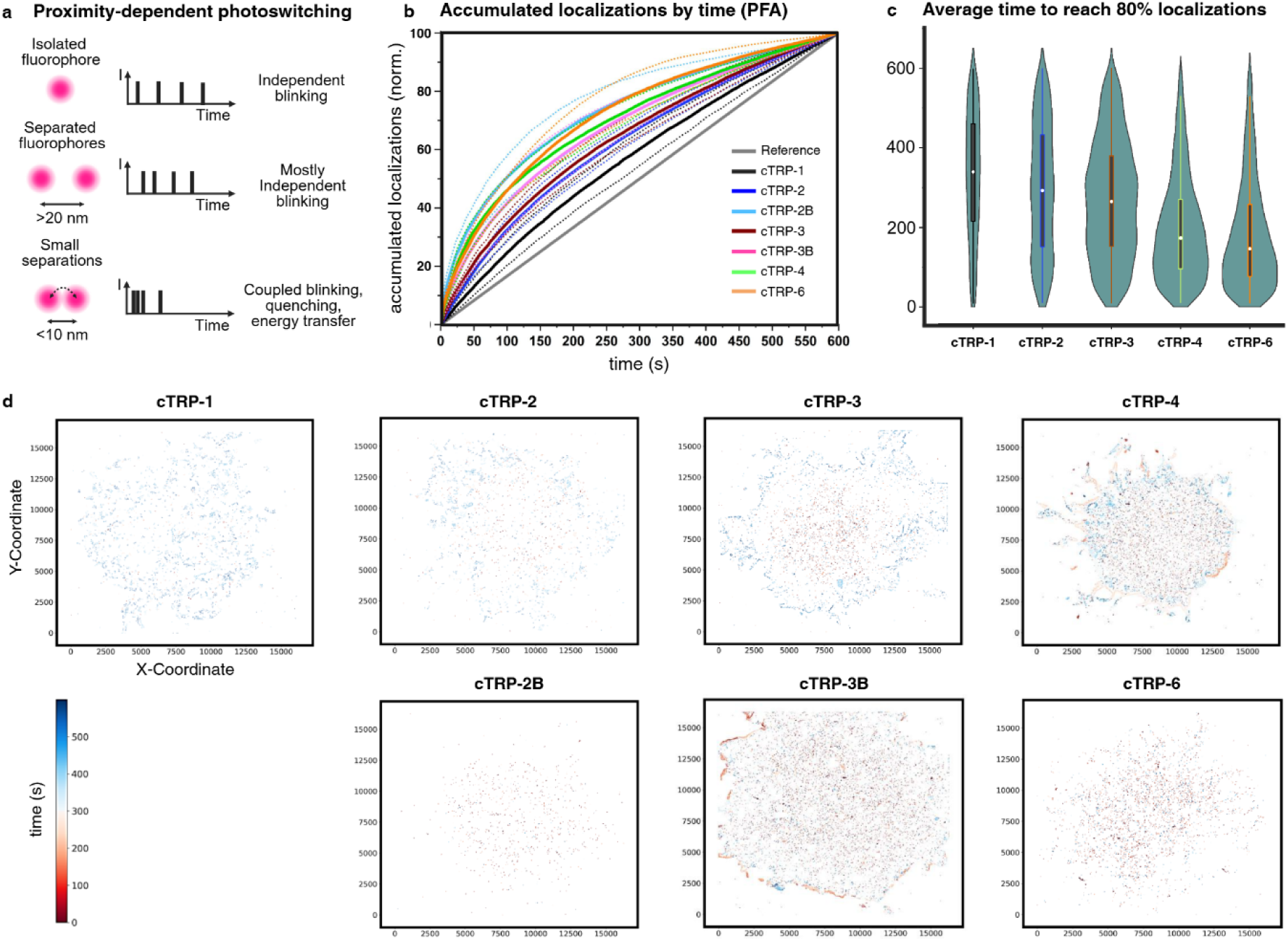
PFA reveals geometry-dependent photoswitching fingerprints. a, Concept of proximity-dependent photoswitching: closely spaced fluorophores can undergo coupled blinking, quenching or energy transfer. **b,** Normalized accumulation of localizations over time for Cy5-labeled cTRP PicoRulers. **c,** Average time required to reach 80% of accumulated localizations for each cTRP geometry. **d,** Spatial maps of the time required to accumulate 80% of localizations within individual cTRP clusters.

Genetically encoded cTRP variants labeled with Cy5 were imaged under identical switching conditions. We analyzed the localization accumulation kinetics of individual cTRP clusters and compared them with idealized reference traces representing independent fluorophores (Fig. 3b). The resulting photoswitching fingerprints differed across the cTRP panel, indicating that fluorophore arrangement within the protein scaffold affects the temporal accumulation of localizations. In addition to the distributed cTRP-1 to cTRP-6 valency series, compact designs such as cTRP-2B and cTRP-3B provided geometries in which fluorophores were positioned at shorter distances within the same ring architecture. These variants allowed us to assess the contribution of nanoscale spacing independently of fluorophore number.

To quantify these differences, we mapped the time point at which 80% of all localizations within a cluster had accumulated (Figs. 3c,d). These analyses revealed construct-dependent differences in localization accumulation, with higher-valency and more compact fluorophore arrangements showing faster accumulation behavior than single-fluorophore cTRPs. The differences between compact and more widely distributed arrangements of similar valency further indicate that the PFA readout is sensitive not only to fluorophore number, but also to their relative nanoscale spacing.

We further analyzed the underlying blinking properties of the Cy5-labeled cTRPs (Extended Data Fig. 5a). Across the construct series, on-times remained broadly similar, whereas off-times shifted towards shorter values for higher-valency and compact designs. Localization intensities remained broadly comparable across the panel, indicating that the observed PFA differences were not explained by a systematic change in single-event brightness. Instead, the altered off-times and localization accumulation kinetics support proximity-dependent changes in photoswitching behavior. By varying fluorophore number and spacing within a compact protein scaffold, cTRPs thus provide defined test samples for evaluating how nanoscale geometry affects localization probability, blinking kinetics and PFA-based molecular readouts.

### 2.4 DNA-PAINT visualizes valency-dependent organization within compact cTRP scaffolds

Having established programmable labeling within the cTRP scaffold, we next addressed the central challenge of the platform: whether these approximately 10-nm protein rings could be resolved and quantitatively read out as individual nanoscale objects. In particular, we asked whether DNA-PAINT could recover the programmed increase in docking-site valency across cTRPs carrying one to six labeling sites. For this purpose, cTRP variants carrying one, two, three, four or six Tet2Et sites were conjugated to trans-cyclooctene-modified DNA-PAINT docking strands and immobilized on a single-molecule surface (Fig. 4a). Transient binding of Cy3B-labeled imager strands enabled localization-based detection of individual cTRP particles under identical imaging conditions.

**Figure 4.**
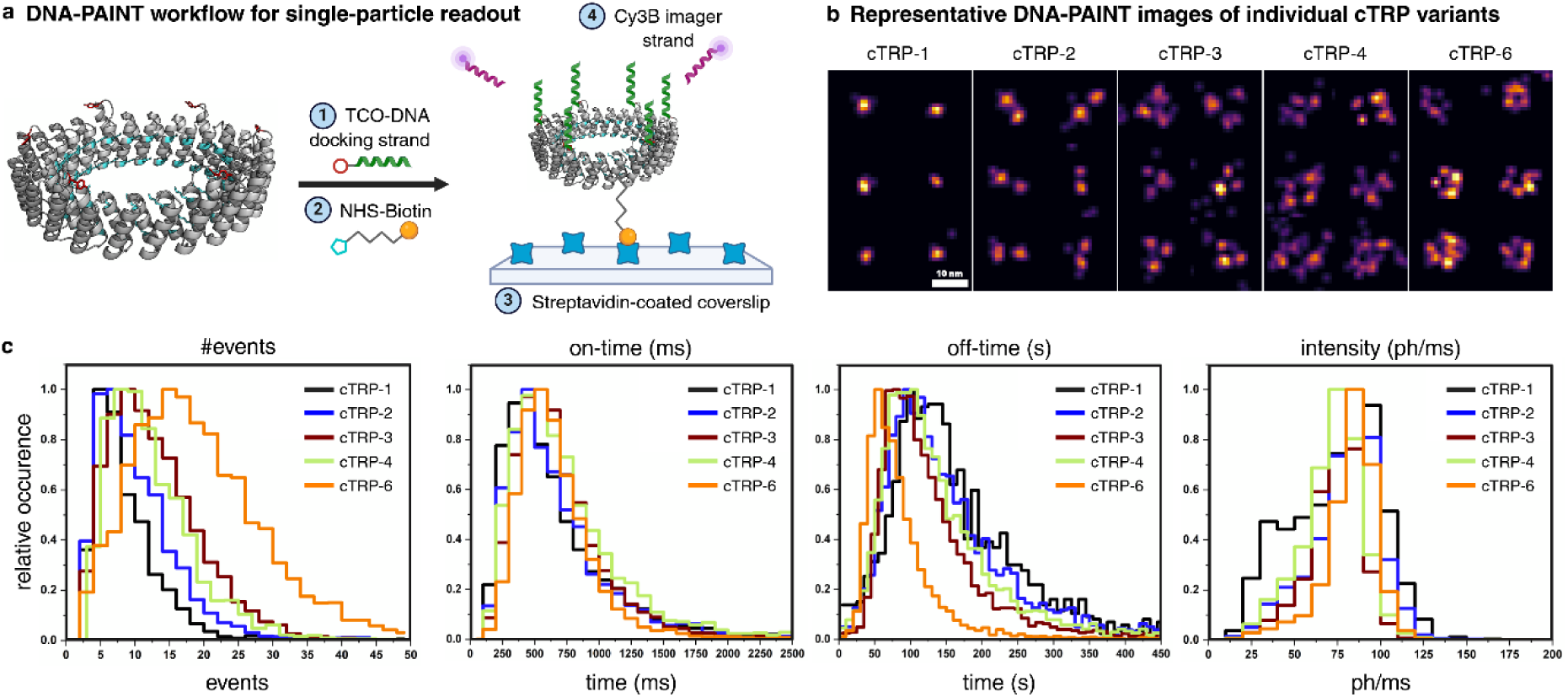
DNA-PAINT visualizes valency-dependent docking-site organization in cTRP PicoRulers. a, DNA-PAINT workflow for single-particle analysis. Tet2Et-containing cTRPs were conjugated to trans-cyclooctene-modified docking strands, biotinylated, immobilized on streptavidin-coated coverslips and imaged with Cy3B-labeled imager strands. **b,** Representative DNA-PAINT images of individual cTRP-1, cTRP-2, cTRP-3, cTRP-4 and cTRP-6 particles acquired under identical conditions. Scale bars, 10 nm. **c,** Distributions of detected binding events, imager-strand on-times, apparent off-times and localization intensities.

Strikingly, DNA-PAINT resolved the programmed valency series within the ∼10-nm cTRP scaffold (Fig. 4b). Individual particles displayed spatially distinguishable binding sites, closely matching the number and arrangement of docking positions encoded in each construct. Thus, the molecular architectures programmed into the protein scaffold could be recovered directly from single-particle fluorescence images. Resolving multiple distinct binding sites within a single approximately 10-nm protein ring establishes cTRP PicoRulers as optically readable molecular standards at the scale of individual protein complexes.

Quantitative analysis of DNA-PAINT binding kinetics further supported the programmed increase in accessible docking-site valency (Fig. 4c, Extended Data Fig. 5b). Under identical imager-strand conditions, the mean number of detected binding events increased overall across the cTRP panel, with cTRP-6 showing the highest event number, whereas on-times and localization intensities remained broadly comparable between constructs. Apparent off-times were also shortest for cTRP-6, consistent with an increased probability of imager-strand binding to particles carrying multiple accessible docking sites. By contrast, the broadly comparable on-times and localization intensities indicate that the differences in event frequency were not driven by major changes in single-event dwell time or brightness.

Together, these experiments establish DNA-PAINT as an orthogonal single-particle assay for probing cTRP PicoRuler valency and docking-site accessibility. They confirm that the engineered protein scaffolds can be functionalized with DNA docking strands, immobilized as single particles and detected as compact nanoscale reference structures.

### 2.5 cTRP PicoRulers are compatible with expansion microscopy

We finally asked whether cTRP PicoRulers could serve as reference structures beyond classical single-molecule imaging. ExM provides a complementary route to molecular-scale fluorescence imaging by physically separating labeled biomolecules before optical readout, but quantitative interpretation requires standards that can report retention, distortion and effective post-expansion spacing. Because cTRPs are compact protein scaffolds with defined click-site positions, we reasoned that they could provide molecular reference structures for evaluating expansion-based workflows, which are mostly incompatible with DNA nanotechnology such as DNA-origami.

To prove this, we used recombinant cTRP-6 PicoRulers carrying six clickable sites (*p*-Azido-L-phenylalanine, pAzF) distributed around the protein ring with a designed nearest-neighbor spacing of approximately 4.5 nm. The proteins were anchored with AcX, expanded approximately eight- to tenfold using a TREx-based workflow, subjected to trypsin digestion and labeled after expansion with DBCO-Alexa Fluor 647 before SRRF imaging (Fig. 5a).^5,24^ Post-expansion click labeling produced readily detectable fluorescence, indicating that cTRP material and chemically accessible click sites were retained through anchoring, digestion, and expansion (Fig. 5c). Individual cTRP particles remained detectable as compact fluorescent structures, supporting their compatibility with expansion-based sample preparation and post-expansion optical readout.

**Figure 5.**
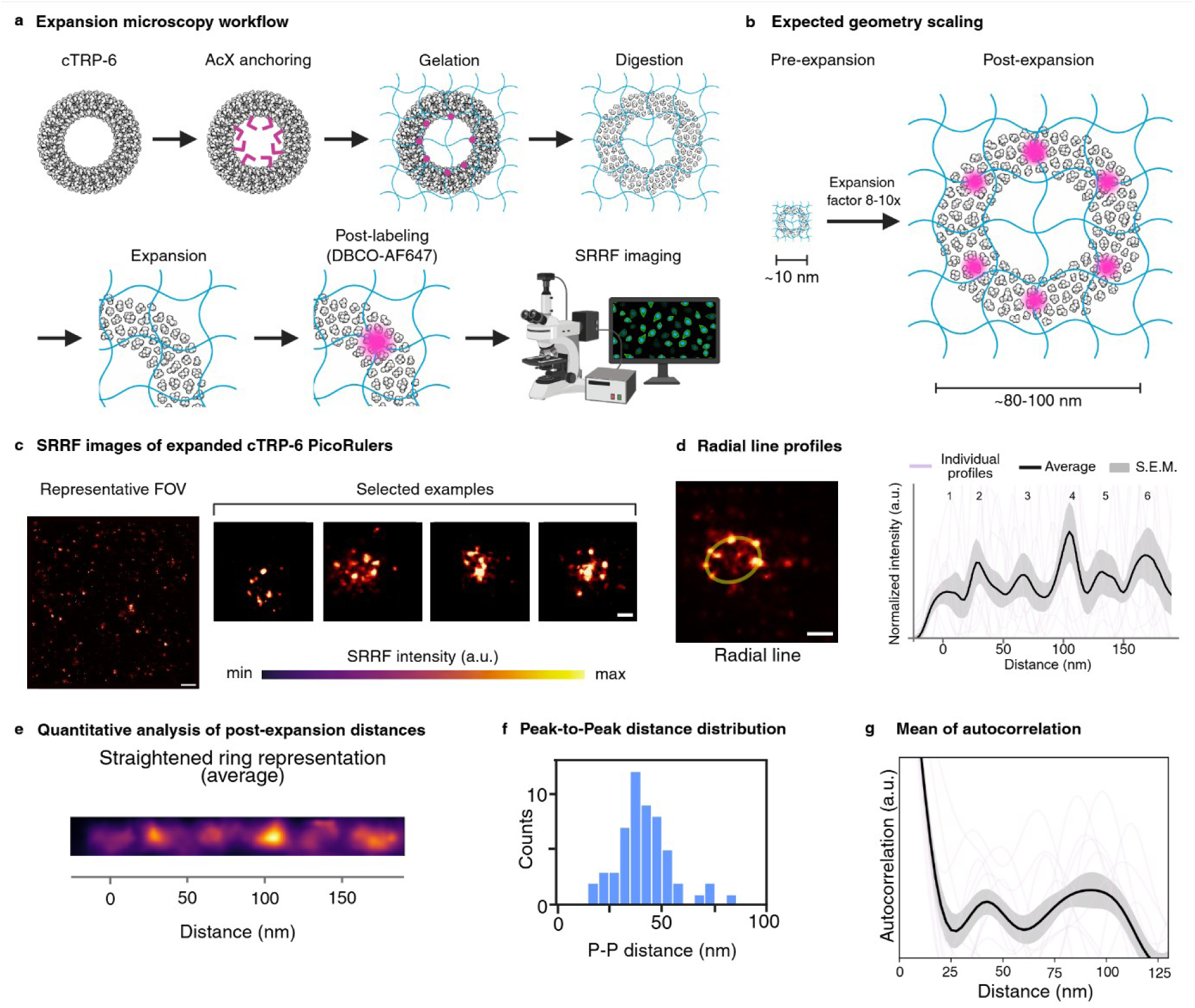
cTRP PicoRulers are compatible with expansion-based imaging workflows. a, TREx-based expansion microscopy workflow for cTRP PicoRulers, comprising AcX anchoring, gelation, digestion and eight- to tenfold expansion, followed by post-expansion labeling with DBCO–Alexa Fluor 647 and SRRF imaging. **b,** Expected scaling of an approximately 10-nm cTRP ring to an apparent diameter of approximately 80–100 nm after expansion. **c,** Representative field of view and selected SRRF reconstructions of expanded cTRP-6 particles; the intensity lookup table applies to selected particles. **d,** Circumferential intensity profiles from ring-like particles. Individual profiles, the average and s.e.m. are shown. **e,** Averaged straightened representation of aligned ring-like particles. **f,** Distribution of adjacent peak-to-peak distances extracted from individual profiles; mean distance, 40.5 ± 1.7 nm. **g,** Autocorrelation profiles of individual particles and their mean ± s.e.m., showing recurring spatial organization after expansion. Scale bars, 1µm (c, FOV) and 50 nm (c, selected examples).

SRRF reconstructions revealed ring-like particles with recurrent circumferential intensity maxima (Fig. 5c). The designed inter-fluorophore spacing of 4.5 nm predicts an apparent separation of 36– 45 nm after eight- to tenfold expansion (Fig. 5b). Intensity profiles traced along the circumference of individual particles showed recurrent peaks, and analysis of adjacent peak positions yielded a mean peak-to-peak distance of 40.5 ± 1.7 nm, closely matching the expected range (Fig. 5d–f). Averaged autocorrelation analysis provided an alignment-independent measure of periodicity and independently supported recurring spatial organization on the expected length scale (Fig. 5g).

These data show that cTRP PicoRulers retain optically accessible nanoscale organization after anchoring, expansion, proteolysis and post-expansion labeling. By confining a defined click site number within an approximately 10-nm protein ring before expansion, cTRP PicoRuler provide a molecular reference object for evaluating signal retention, local expansion and geometric distortion at the scale of individual protein assemblies. cTRP PicoRulers therefore extend programmable protein standards to expansion microscopy and provide a route to benchmarking molecular-scale performance in expanded samples.

## 3. Discussion

Fluorescence microscopy is increasingly being applied at length scales where labeling and fluorophore behavior become major determinants of quantitative accuracy. At distances below 10 nm, stoichiometry, linkage error, accessibility and short-range photophysical interactions can substantially alter apparent molecular organization. Here we introduce cTRP PicoRulers as programmable protein reference standards in which fluorophore number and spacing can be varied within the same approximately 10-nm scaffold.

The central advance of the cTRP platform is the combination of geometric programmability with a protein-based labeling environment. DNA origami provides highly defined synthetic geometries, whereas cellular structures such as nuclear pores offer biological context but cannot be freely redesigned. cTRPs bridge these reference classes and extend previous protein-based standards such as PCNA PicoRulers beyond a fixed trimeric architecture to a modular series of fluorophore configurations. DNA-PAINT resolved the programmed valency series at the single-particle level, demonstrating that the molecular designs remain optically accessible within the compact scaffold. At the same time, the short inter-site distances make cTRPs promising benchmark samples for high-precision localization methods such as MINFLUX and RESI, enabling validation of molecular distance measurements in the nanometer regime.

Beyond optical resolution, cTRPs provide a means to benchmark proximity-dependent fluorophore behavior. The geometry-dependent molecular fingerprints observed by PFA show that closely spaced fluorophores cannot always be treated as independent emitters. By controlling valency and spacing within the same scaffold, cTRP PicoRulers convert such interactions from an uncontrolled source of bias into a measurable property of the reference sample.

The platform can also be deployed in biological and expansion-based imaging environments. Recombinant tethering and genetically encoded membrane display place the standards at the cell surface, where crowding, orientation and intermolecular proximity influence fluorescence readouts. cTRP particles also retained measurable periodic organization after anchoring, proteolytic expansion and post-expansion labeling, indicating that the scaffold can serve as a molecular reference object for evaluating retention and local expansion at the scale of individual protein assemblies.

Together, cTRP PicoRulers provide programmable protein standards for molecular-scale fluorescence microscopy. Their compact, genetically encoded and chemically addressable architecture should facilitate the development of reference libraries tailored to different fluorophores, geometries and imaging workflows.

## 4. Methods

### Expression and Suppressor Plasmids

Plasmids encoding wild-type cTRP 1–4 (pET15HE backbone) were kindly provided by Barry L. Stoddard.^19^ The plasmids for expressing clickable self-assembling cTRP 1-6 were generated via site-directed mutagenesis and cloning into the wild-type plasmids. The genes encoding clickable cTRP 1-6 as a single polypeptide chain were synthesized commercially (Genscript) and cloned into pBAD expression plasmids. The suppressor plasmids pAJE3-E7 for Tet2Et incorporation and pEVOL-pAzF-2t1 for pAzF incorporation were gifts from Ryan Mehl (Addgene plasmid #214359)^20^ and Farren Isaacs (Addgene plasmid #73546),^25^ respectively. GCExpress suppressor plasmid was used for the incorporation of TCO*A in mammalian cells.^26^

### Expression of Wild-Type cTRPs

Wild-type cTRPs (1–4) were expressed as previously described,^19^ with modifications. Briefly, plasmids were transformed into *E. coli* C41(DE3) cells (Sigma-Aldrich) and plated on 2×YT (Melford) agar with 100 μg mL^-1^ ampicillin (Melford). A single colony was used to inoculate an overnight culture at 37 °C, 200 rpm. The next day, the culture was diluted 1:100 into fresh medium and grown to OD₆₀₀ of 0.8. Cells were chilled on ice for 10 min, then induced with 0.3 mM IPTG (Carl Roth) and incubated at 16 °C for 24 h. Cells were harvested by centrifugation and stored at −20 °C.

### Expression of Clickable cTRPs

Self-assembling cTRP 1-6 were expressed similarly to their wild-type counterparts with adjustments. Expression (pET15HE) and suppressor plasmids were co-transformed into *E. coli* C41(DE3) cells and maintained with ampicillin and 50 μg mL^-1^ spectinomycin (Sigma-Aldrich). At OD₆₀₀ of 0.8, the medium was replaced with BactoMedia (Thermo Fisher) containing 0.75% glycerol and both antibiotics. Tet2Et (GCE4All, Oregon State University) was added at 0.5 mM final concentration, and after 30 min, protein expression was induced with 0.3 mM IPTG at 30 °C. Single-chain cTRP 1-6 were similarly expressed in C321.ΔA.opt cells (Addgene, #87359) at 30 °C. Expression (pBAD) and suppressor plasmids were co-transformed and maintained with ampicillin and spectinomycin. Protein expression was induced with 0.1% L-Arabinose. For expression of pAzF-containing cTRPs, pAzF (Lumiprobe) was added at 1 mM final concentration, and after 30 min protein expression was induced by 0.2% L-Arabinose.

### Purification of Wild-Type cTRPs

Cell pellet containing single-chain cTRP (1) was resuspended in lysis buffer composed of 25 mM PBS pH 8.0 (Carl Roth), 150 mM NaCl (Carl Roth), 20 mM imidazole (Carl Roth), protease inhibitor cocktails (Sigma-Aldrich), DNase (AppliChem), lysozyme (Carl Roth), and BugBuster 10X (Merck Millipore). After 20 min, lysates were clarified by centrifugation and the supernatant was purified on a HisTrap HP column (Cytiva, 1 mL) using an ÄKTA Pure 25 M system (Cytiva). Elution was performed with 25 mM PBS pH 8.0, 150 mM NaCl, 300 mM imidazole. Fractions containing cTRP (1) were concentrated and further purified by size-exclusion chromatography (SEC) on a HiLoad Superdex 200 PG 16/600 column (Cytiva) using 20 mM HEPES pH 8.0 (Carl Roth), 150 mM NaCl.

Self-assembling cTRPs (cTRP 2-4) were lysed and initially purified as described above, followed by ion-exchange chromatography (IEX) using a MonoQ 10/100 GL column (Cytiva, 8 mL) in 25 mM Tris pH 7.5 (Carl Roth) with a linear gradient of 200 mM – 1 M NaCl (0 – 50% B over 10 column volumes). A final SEC step in 20 mM HEPES pH 8.0, 150 mM NaCl completed the purification.

### Purification of Clickable cTRPs

Single-chain clickable cTRPs (cTRP 1-6) were purified using the same protocol as wild-type cTRP (1), except that SEC was performed in 25 mM Tris pH 7.5, 1 M NaCl.

### Characterization of cTRPs

All wild-type and clickable cTRP constructs were analyzed by SDS-PAGE under reducing and non-reducing conditions. To demonstrate that the lower position of wild-type cTRP-1 band was due to the open conformation of the cTRP ring, the protein was crosslinked with 2400 equivalents of Bis(NHS)PEG-9 (Thermo Fisher) before SDS PAGE. Additionally, to show that cTRP rings were formed through the assembly of smaller subunits linked by disulfide bridges, wild-type cTRP 2-4 were partially reduced with 100 mM DTT in 100 mM Tris pH 7.5 for 24 h. DTT was then removed using a Zeba spin desalting column (Thermo Fisher, 7K MWCO), and the samples were analyzed by SDS-PAGE under non-reducing conditions. Transmission Electron Microscopy (TEM) was also performed on wild-type cTRP (1) to confirm ring formation.

### Transmission Electron Microscopy (TEM) and Analysis

Ring formation was validated by TEM (JEM 1011, JEOL) using negative staining. Carbon-coated grids (300 mesh) were glow-discharged before use, incubated with 20 μL of 500 nM cTRP solution for 2 min, then blotted with filter paper. The grids were dipped three times into a 0.75% uranyl acetate (EMS) and blotted, followed by a final staining step (0.75% uranyl acetate, 45 s), blotting and air drying for 24 h. TEM images were analyzed using ImageJ (v2.3.0/1.53t).

### Site-specific Labeling of cTRPs with Dye/Docking Strand

Clickable cTRPs (cTRP 1-6) were reacted with either 20 equivalents of sCy5-TCO4 (BroadPharm) or P1-TCO4 docking strand (Biomers) relative to Tet2Et in 25 mM Tris pH 7.5, 1 M NaCl for 24 hours. Labeled proteins were purified by SEC on a Superdex 200 Increase 10/300 GL column (Cytiva) in 20 mM HEPES pH 8.0, 150 mM NaCl. Fractions containing labeled cTRPs were pooled and concentrated to 100 μL.

### Determination of Degree of Labeling (DOL)

Protein concentrations of cTRP constructs (cTRP-1 to cTRP-6) were determined by intrinsic fluorescence because their low aromatic amino acid content precluded reliable quantification by absorbance at 280 nm. Variant-specific calibration curves were generated from unlabeled proteins of known concentration and used to quantify the corresponding labeled samples under identical measurement conditions. Fluorophore concentrations were determined independently by UV/Vis absorption spectroscopy using the characteristic absorbance maximum and molar extinction coefficient of the respective dye. The DOL was calculated as the molar ratio of fluorophore to protein.

### Functionalization of cTRPs with Biotin/Halotag ligand

Dye/docking strand prelabeled cTRPs (cTRP 1-6) were reacted with either 20 equivalents of NHS-Biotin (Sigma Aldrich) or 100 equivalents of NHS-Halotag ligand (BLD Pharm) relative to cTRPs in 100 mM HEPES pH 8.3, 150 mM NaCl for 1 hour. The reaction was quenched by addition of 1 M Tris pH 8.0 for 15 min before purification by SEC on a Superdex 200 Increase 10/300 GL column (Cytiva) using 20 mM HEPES pH 8.0, 150 mM NaCl as elution buffer. Fractions containing functionalized cTRPs were pooled and concentrated to 100 μL.

### Tethering of Recombinant cTRP PicoRulers on the Plasma Membrane via HaloTag

An extracellular HaloTag7 construct was generated using the pDisplay mammalian expression vector (Invitrogen, Thermo Fisher Scientific, Cat. No. V66020), with HaloTag7 positioned between the N-terminal Igκ leader sequence and the C-terminal myc epitope–PDGFR transmembrane anchor, as previously described by Li *et al.*^27^ HEK293T/COS-7 cells were transfected with this construct to display HaloTag7 proteins on the extracellular surface of the plasma membrane. HaloTag ligand-functionalized cTRP PicoRulers were incubated with cells for 15 min at room temperature to enable covalent tethering to extracellular HaloTag. Cells were then washed 3 times with imaging buffer to remove unbound cTRP PicoRulers before confocal microscopy.

### Preparation of Genetically Encoded Membrane-Displayed cTRPs

cTRP 1–6 constructs were engineered with an N-terminal signaling peptide, His6X tag, FLAG tag, and thrombin cleavage site, as well as a C-terminal (EAAAK)₆ rigid linker, HRV3C cleavage site, and (G₄S)₃ flexible linker, followed by fusion to GPI-anchored proteins for plasma membrane display. As the NXT motif represents a consensus sequence for N-linked glycosylation in mammalian cells, the GSNXT insertion motif used for bacterial expression was replaced with GSGXL (GXL = GGG TAG CTC), an optimized codon context for amber suppression in mammalian cells.^23^ The corresponding genes were commercially synthesized (GenScript) and cloned into pcDNA3 vectors. HEK293T/COS-7 cells were co-transfected with the respective expression plasmids and the GCExpress system and cultured for 24 h in the presence of 250 µM TCO*A prior to labeling. Membrane-displayed clickable cTRPs were site-specifically labeled through bioorthogonal click chemistry. Excess fluorophore was removed by washing with imaging buffer prior to confocal microscopy.

### DNA-PAINT Single-Molecule Surface Preparation

The surfaces of 8-well chambered cover glass with high performance cover glass (Cellvis, C8-1.5H-N) were washed once with PBS (Sigma-Aldrich) and treated with 2% Hellmanex (Hellma) for 1 h. Afterwards, the treated chambers were washed 3× with PBS, incubated with 1 M KOH (Fluka) for 20 min, and rewashed with PBS. Thereafter, the surfaces were incubated with 10% polyethylene glycol 400 (Fluka) overnight at 4 °C, followed by washing 3x with PBS. The surfaces were incubated with 0.5 g L^−1^ BSA-Biotin (ThermoFisher) in PBS overnight at 4 °C. On the next day, the chambers were washed 3× with PBS before incubation with 0.5 g L^−1^ Neutravidin (ThermoFisher) in PBS for 20 min. Afterwards, the treated chambers were washed 2× with PBS. cTRP samples were diluted to ≈1 nM in PBS. Briefly, the cleaned surfaces were incubated with the diluted protein samples for ≈10 s and washed 3× with PBS. The samples then were washed and covered in PBS-based imaging buffer containing 5 mM Tris, 1 mM EDTA and 10 mM MgCl_2_ adjusted to pH 7.6.

### Super-Resolution Imaging and Analysis

All measurements were performed on an inverted wide-field fluorescence microscope (IX-71, Olympus). For excitation of Cy5 a 641-nm diode laser (Cube 640-100 C, Coherent) in combination with a clean-up filter (Laser Clean-up filter 640/10, Chroma) was used. For excitation of Cy3B imager strands (DNA-PAINT), a 561-nm diode laser (Genesis MX561-500 STM, Coherent) with irradiation intensity of ≈0.25 – ≈1 kW cm^−2^ in combination with a clean-up filter (Laser Clean-up filter 561/14, Chroma) was used. All measurements were performed using circular polarized light by mounting a quarter-wave plate (Thorlabs,) within the excitation path. The laser beam was focused onto the back focal plane of the oil-immersion objective (×60, NA 1.45, Olympus). For measurements, emission light was separated from the illumination light using a dichroic mirror (HC 560/659 (Cy5) or FF580-FDi01 (DNA-PAINT), Semrock) and spectrally filtered by a bandpass filter (FF01-679/41-25 (Cy5) or BrightLineHC-600/50 (DNA-PAINT), Semrock). Images were recorded with an electron-multiplying CCD camera chip (iXon DU-897, Andor). Pixel size for data analysis was measured to 128 nm. For PFA measurement 120 000 images with an exposure time of 5 ms (frame rate 200 Hz) and irradiation intensity of roughly ≈3.5 kW cm^−2^ were recorded using highly inclined and laminated optical sheet (HILO) illumination. All PFA experiments were performed in PBS-based photoswitching buffer containing 100 mM β-mercaptoethylamine (Sigma-Aldrich) adjusted to pH 7.6. For DNA-PAINT measurement, 18 000 images with an exposure time of 100 ms (frame rate 10 Hz) were recorded by total internal reflection fluorescence microscopy illumination. DNA-PAINT experiments were performed with 3 nM imager strand concentration (5′–3′: CTA GAT GTA T, Eurofins), 3′-modified with Cy3B, in PBS-based imaging buffer containing 5 mM Tris, 1 mM EDTA, and 10 mM MgCl_2_ adjusted to pH 7.6. All SMLM results were analyzed with rapidSTORM3.3 and the highly resolved DNA-PAINT cTRP pictures were reconstructed with ThunderSTORM.^28,29^ For analyzing the blinking kinetics, fluorescent spots containing more than 115 (PFA)/1000 (DNA-PAINT) photons per frame were analyzed. The estimation of the number of localizations per fluorophore was calculated by using the tracking function (Kalman filter) of rapidSTORM3.3. Fluorescent spots were tracked over the whole image stack within a tracking radius of 200 nm and exported as tracked localization file. A custom written python script was used to calculate the number of frames between on-time events of the same fluorescent spot within the defined tracking radius (off-time) as well as the number of on-time events per tracked spot.

### PFA of Localization Kinetics and Cluster Maturation

Localization data obtained from rapidSTORM3.3 were further analyzed using a custom-written Python script (Python 3.6). Localizations were filtered based on intensity thresholds to exclude low-confidence events. Spatial clustering was performed using the DBSCAN algorithm with a user-defined clustering radius (10 nm) and minimum number of localizations per cluster (5 locs.). For each identified cluster, localization events were temporally ordered based on frame number and converted into physical time using the acquisition exposure time. The total acquisition time was defined by the first and last recorded frame. To quantify cluster maturation dynamics, the time required to reach 80% of all localizations within a cluster was calculated. Clusters were visualized by plotting their spatial coordinates and color-coded according to their 80% time. Summary statistics, including mean 80% times and localization counts per cluster, were extracted for further analysis.

### ExM and SRRF imaging

For expansion microscopy 10-80 µg of purified cTRP-1 6TAG pAzF was transferred into 100 uL crosslinking solution consisting of 150 mM NaHCO_3_ in PBS (pH 8.3), supplemented with 0.3 mg mL^-1^ Acryloyl-X (A-20770, Thermo Fisher Scientific). cTRPs in crosslinking solution were transferred onto 12 mm coverslips, placed in a humidified chamber and incubated overnight at 4°C. After incubation, samples were dried under a fume hood, until only a thin liquid film remained. Importantly, the drying process was stopped before visible salt precipitation occured. Immediately after semi-drying, samples were flipped onto a 50 µL drop TREx monomer consisting of 1.1 M sodium acrylate, 2.0 M acrylamide, 90 ppm N,N’-methylenebisacrylamide, 1.5 ppt APS, and 1.5 ppt TEMED in 1x PBS, and incubated for 5 min on ice. Gelation proceeded for an additional 1.5 h at 37°C in a humidified chamber. TREx samples were homogenized either by proteolytic cleavage using Trypsin overnight at 37°C (0.015% Trypsin-EDTA in 35 mM Tris, 0.7 mM EDTA, 0.35% Triton-X-100 and 0.5 M NaCl adjusted to pH 8.0). Gels were washed at least three times in PBS, then stained with 30 µM DBCO-AF647, or DBCO-sulfo-Cy5 overnight. Gels were washed again at least three times in PBS, then expanded in Milli-Q water. Fully expanded samples were cut to size then mounted on APTES- or PDL-coated 1-well chambered slides (Cellvis, #C1-1.5H-N) for SRRF imaging.

SRRF imaging was performed using a Leica TCS SP8 confocal laser scanning microscope equipped with a 63x/1.45 NA oil immersion objective. Cy5 or Alexa Fluor 647 was excited using the 633-nm laser line, with appropriate emission detection settings selected according to the LAS X microscope control software. Images of 128 x 128 or 256 x 256 pixels were acquired on a HyD detector in photon counting mode with 8-bit pixel depth and a theoretical pixel size set to 50 nm. For each position, a time-series of 2500 frames was acquired with a frequency of 13 – 25 Hz. Movies of fluorescence fluctuations were processed using the ONE microscopy plugin for ImageJ. For SRRF processing, the standard pre-set for the ONE plugin was used with a radiality magnification of 20, and 8 ring axis. The temporal analysis mode was set to temporal radiality auto-correlation (TRAC 4^th^ order).

## Supporting information

cTRP_PicoRulers_Supplementary_Information_v1

## Acknowledgements

M.B. and D.H. contributed equally to this work. The authors thank J. Iff and E. Maier for cell culture support, as well as C. Stigloher and D. Bunsen from the Department of Electron Microscopy, Biocenter, University of Würzburg for their support during quality control of cTRP samples. Figures created with BioRender.com. This project is supported by the Federal Ministry for Economic Affairs and Climate Action (BMWK) based on a decision by the German Bundestag (Grant Agreement No. KK5665801HV4 to G.B.). M.S. acknowledges funding from the European Research Council (ERC) under the European Union’s Horizon 2020 research and innovation programme (grant agreement No 835102). Open access funding enabled and organized by Projekt DEAL.

## Conflict of Interest

The authors declare no conflict of interest.

## Data Availability Statement

All data that support the findings described in this study are available in the manuscript and the related supplementary information, and from the corresponding authors upon reasonable request. Plasmids will be made available via Addgene.

## Extended Data Figure Legends

**Extended Data Fig. 1.**
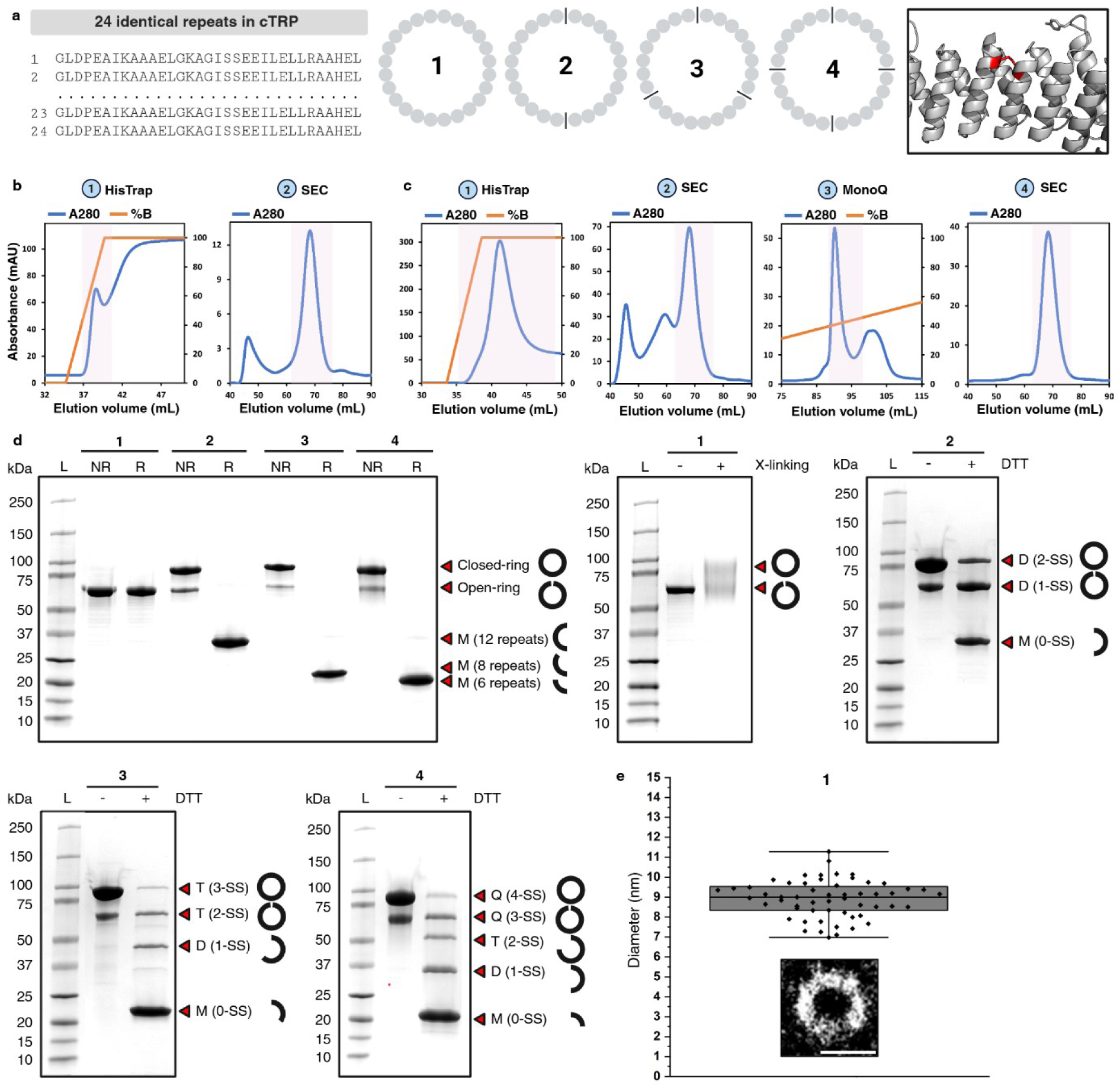
Structural and biochemical validation of single-chain and self-assembling wild-type cTRPs. a, Left, cTRP sequence showing the 24 identical repeats that constitute the scaffold. Middle, cTRP rings can be produced either as a single polypeptide chain (1) or by assembly of smaller subunits linked by disulfide bonds shown as black lines (2–4). Right, an intersubunit disulfide bond is highlighted in red. **b,** Purification workflow for single-chain cTRP (1). **c,** Purification workflow for self-assembling cTRP (4). **d,** Top left, SDS–PAGE analysis under reducing (R) and non-reducing (NR) conditions showing the expected migration patterns of single-chain and self-assembling cTRP constructs. Top middle, crosslinking of single-chain cTRP (1) supports assignment of open- and closed-ring conformations. The remaining three gels show partial reduction of self-assembling cTRPs (2–4), supporting formation of the rings through assembly of smaller disulfide-linked subunits. L, molecular-mass ladder; M, monomer; D, dimer; T, trimer; Q, quartemer; SS, disulfide bridges. **e**, Transmission electron microscopy of purified single-chain cTRP (1) confirming formation of compact ring-shaped particles with an outer diameter of approximately 10 nm. Scale bar, 10 nm.

**Extended Data Fig. 2.**
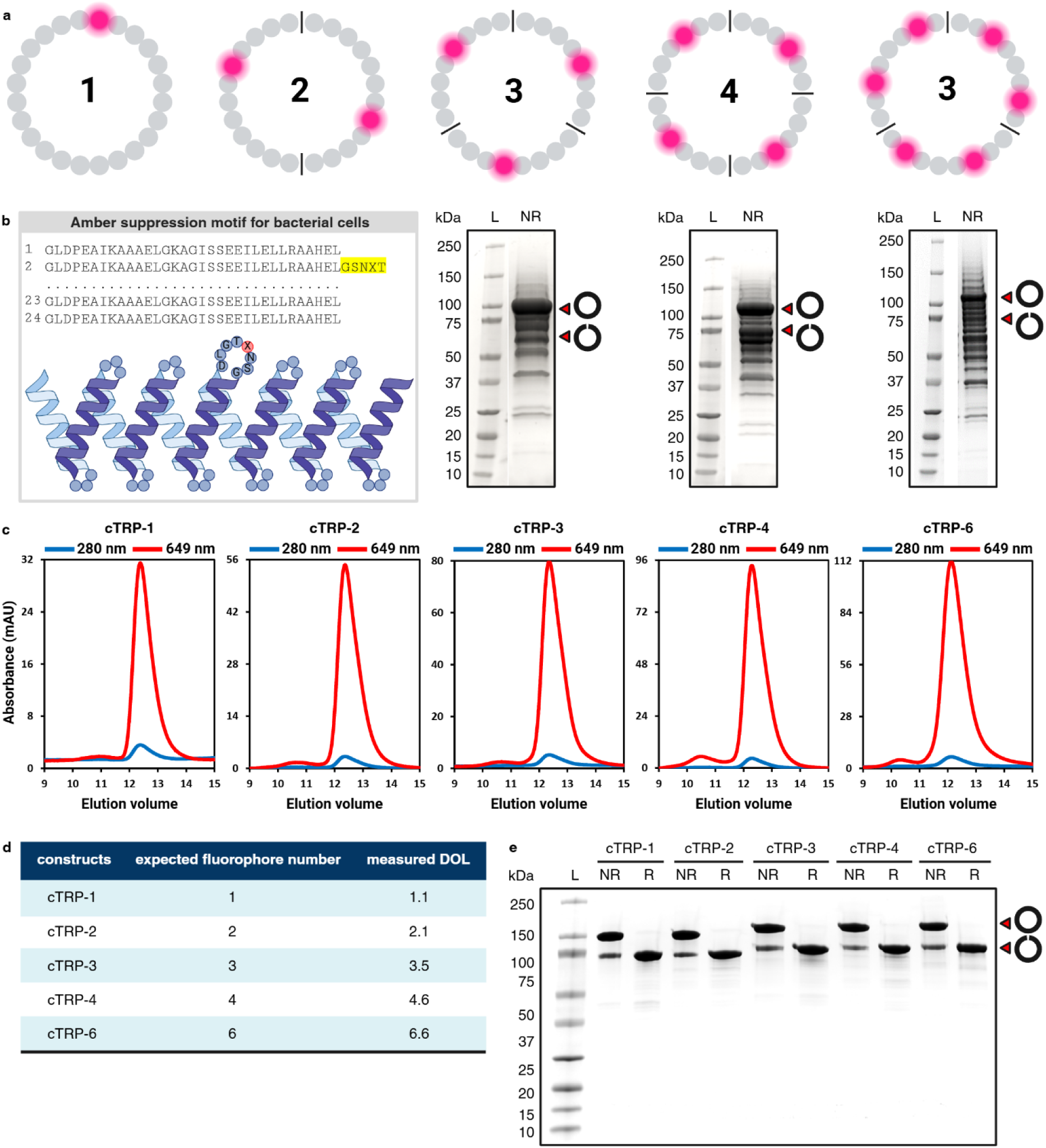
Engineering, production and bioorthogonal labeling of clickable cTRP PicoRulers. **a**, Design of self-assembling cTRP constructs carrying one, two, three, four or six Tet2Et incorporation sites. Black lines indicate intersubunit disulfide bridges. Assembly of subunits amplifies the number of clickable sites, but production becomes increasingly challenging as the number of amber codons increases. Corresponding non-reducing SDS–PAGE analyses show greater product heterogeneity at higher amber-codon numbers, thereby complicating the isolation of fully assembled cTRP rings. **b,** Boxed schematic, insertion of a GSNXT motif into a surface-exposed cTRP loop to promote efficient Tet2Et incorporation by amber suppression. **c,** Size-exclusion chromatography (SEC) profiles of Tet2Et-containing single-chain cTRPs after bioorthogonal labeling with sCy5-TCO, demonstrating fluorescent labeling across the construct panel. **d,** Expected and measured degrees of labeling for cTRP-1 to cTRP-6. **e,** SDS– PAGE analysis of clickable single-chain cTRPs expressed in the RF1-deficient *E. coli* strain C321.ΔA.opt. L, molecular-mass ladder; NR, non-reducing; R, reducing.

**Extended Data Fig. 3.**
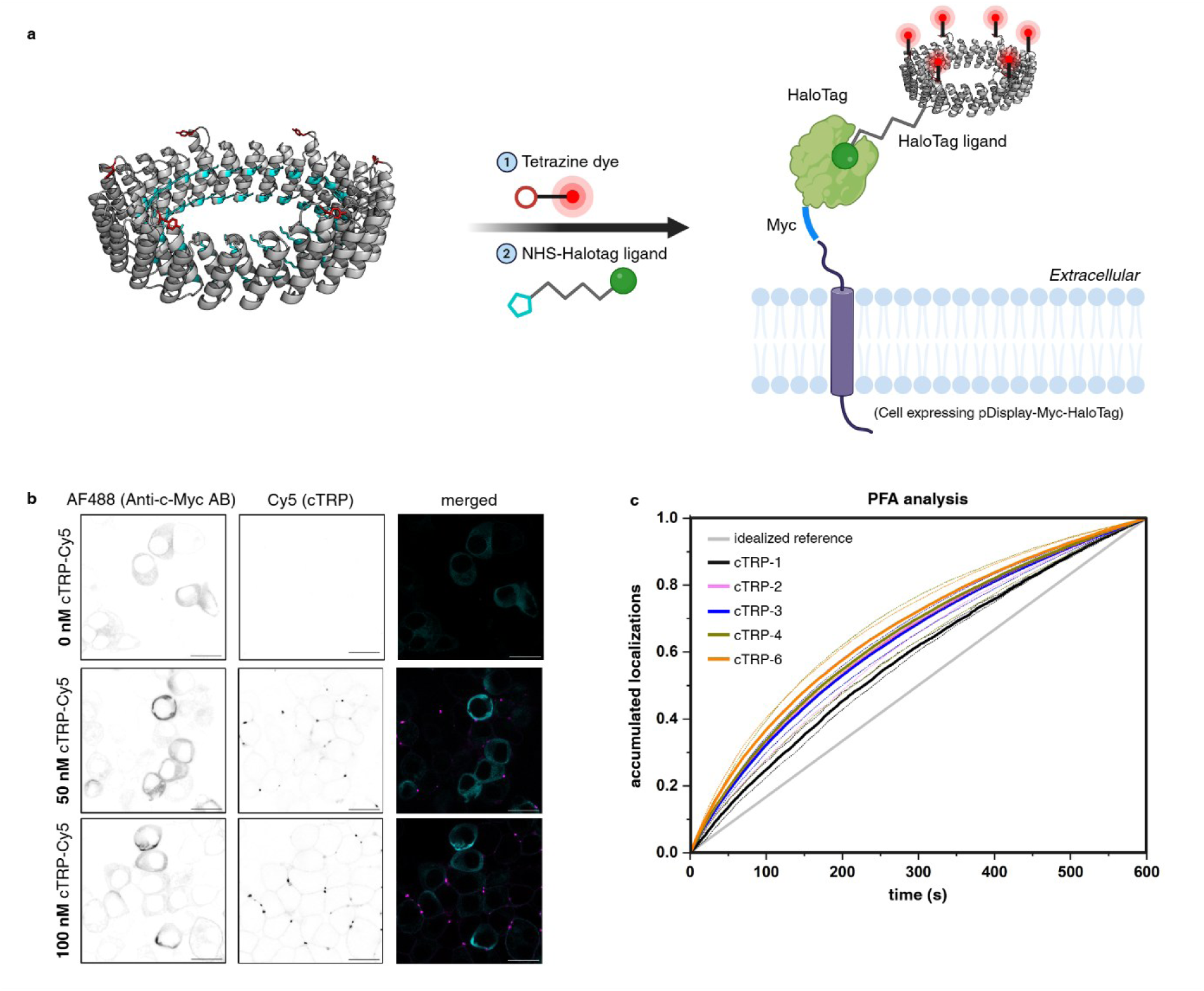
HaloTag-mediated cell-surface tethering and photoswitching analysis of recombinant cTRP PicoRulers. **a**, Schematic of sequential cTRP functionalization with sCy5-TCO and NHS–HaloTag ligand, enabling cell-surface covalent tethering through extracellular HaloTag. **b**, Representative confocal images of cells expressing Myc-tagged extracellular HaloTag after incubation with 0, 50 or 100 nM dual-labeled cTRP PicoRulers. Anti-c-Myc–AF488 marks extracellular HaloTag at the plasma membrane, whereas the Cy5 signal reports cell-surface-tethered cTRP PicoRulers. Merged images show colocalization of Cy5-labeled cTRPs with HaloTag-positive plasma membranes. Scale bars, 20 µm. **c**, Normalized localization-accumulation traces from photoswitching fingerprint analysis of membrane-tethered cTRP-1, cTRP-2, cTRP-3, cTRP-4 and cTRP-6 reveal construct-dependent localization-accumulation profiles.

**Extended Data Fig. 4.**
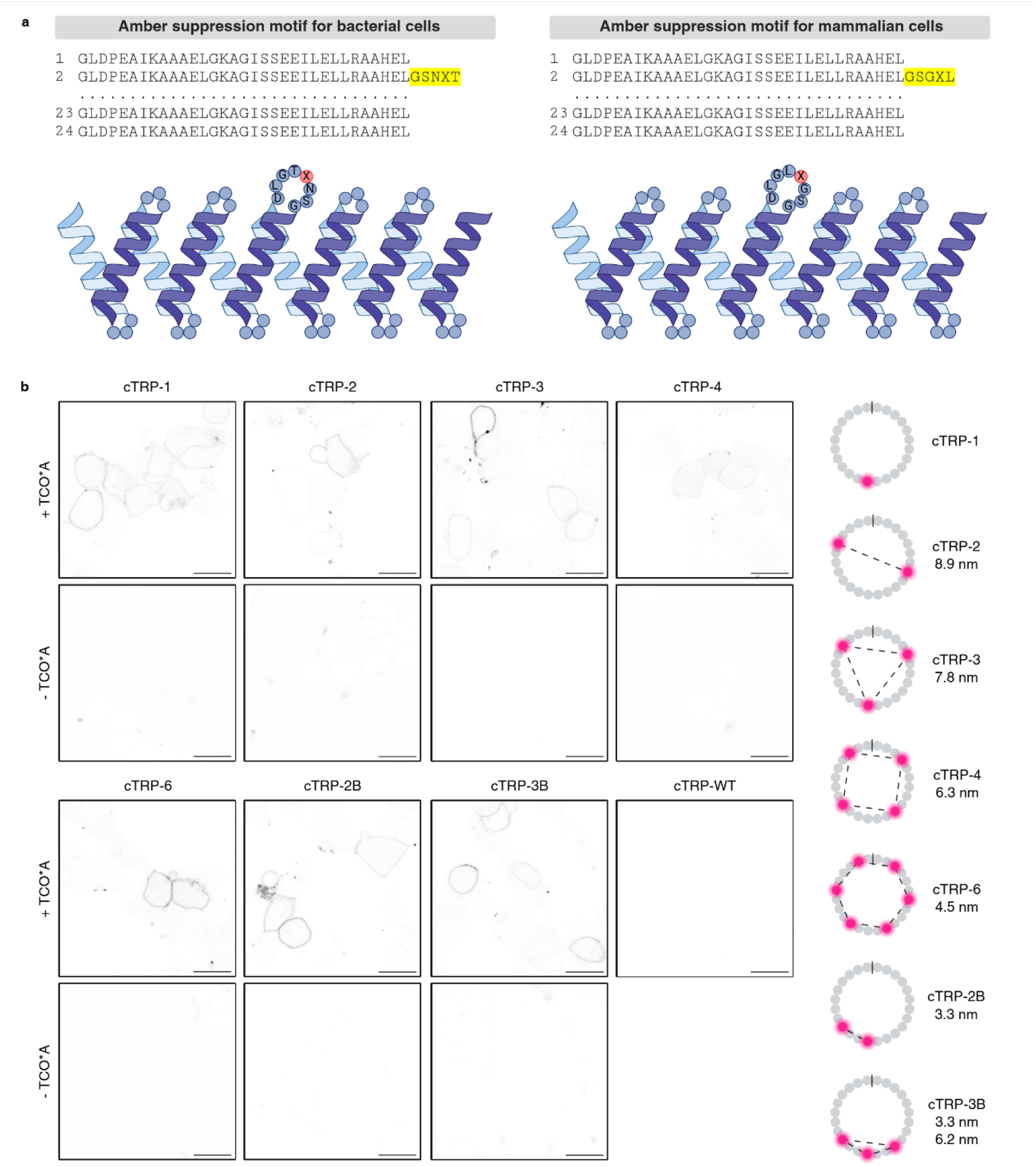
Genetically encoded membrane display of cTRP PicoRulers. a, Replacement of the bacterial GSNXT amber-suppression motif with the mammalian-compatible GSGXL motif to avoid potential N-linked glycosylation while supporting ncAA incorporation in mammalian cells. **b,** Representative confocal images of genetically encoded membrane-displayed cTRP PicoRulers expressed in the presence or absence of TCO*A following bioorthogonal fluorophore labeling. Schematics on the right show the programmed labeling-site arrangements and approximate inter-fluorophore distances in the indicated constructs. Scale bars, 20 µm.

**Extended Data Fig. 5.**
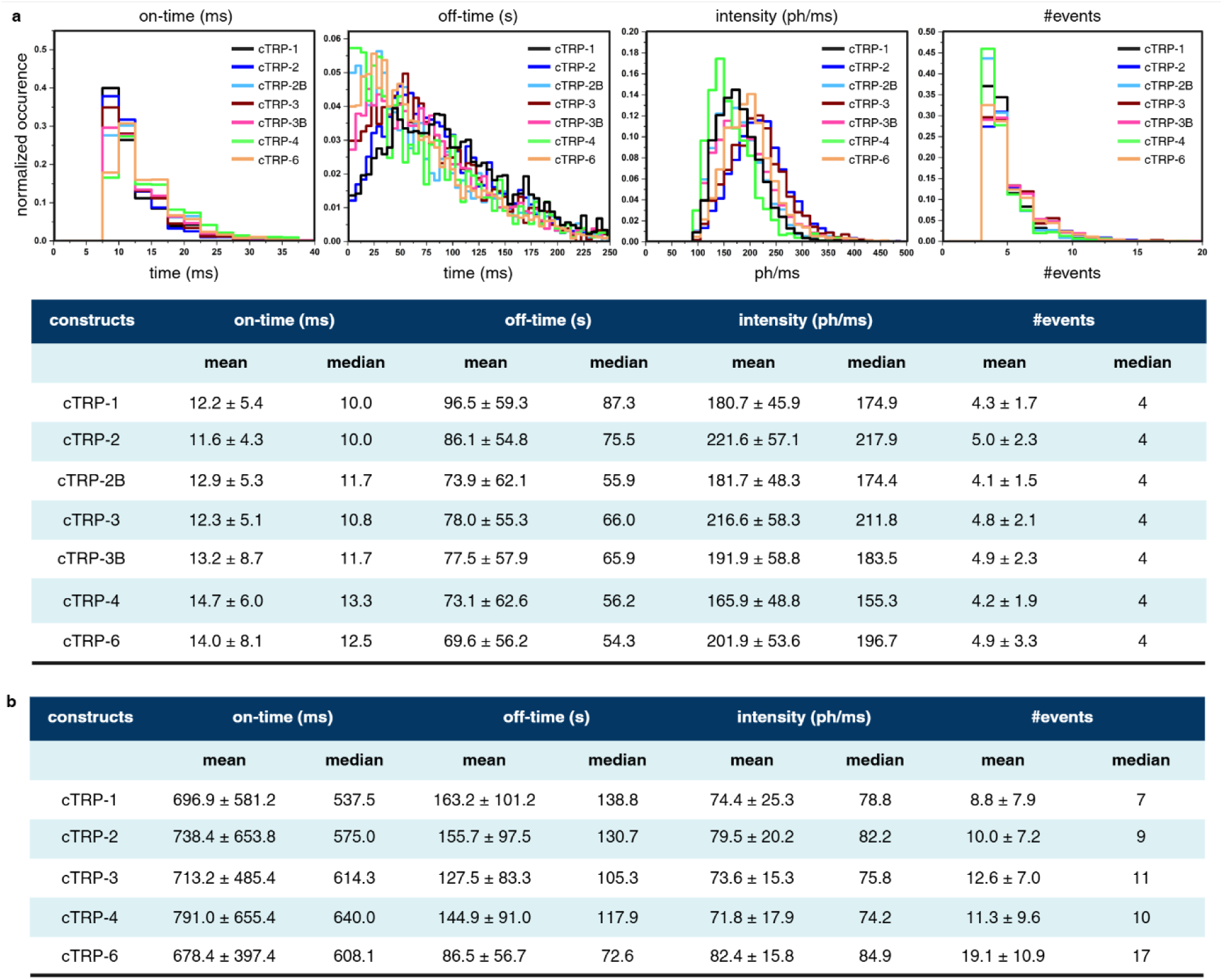
Kinetic analysis of *d*STORM photoswitching and DNA-PAINT binding across cTRP PicoRulers. a, Photoswitching analysis of genetically encoded membrane-displayed cTRP PicoRulers under identical dSTORM conditions. Top, normalized distributions of on-times, off-times, localization intensities and detected event numbers. Bottom, corresponding mean ± s.d. and median values. **b,** Summary of DNA-PAINT kinetic parameters and detected binding events for cTRP-1, cTRP-2, cTRP-3, cTRP-4 and cTRP-6. Mean ± s.d. and median values are shown for imager-strand on-times, apparent off-times, localization intensities and event numbers.

## Notes

### Competing Interest Statement

The authors have declared no competing interest.

## References

1. Reinhardt, S. C. M. et al. Ångström-resolution fluorescence microscopy. Nature 617, 711– 716 (2023).

2. Strauss, S. C Jungmann, R. Up to 100-fold speed-up and multiplexing in optimized DNA-PAINT. Nat. Methods 1–3 (2020) doi:10.1038/s41592-020-0869-x.

3. Sahl, S. J. et al. Direct optical measurement of intramolecular distances with angstrom precision. Science 386, 180–187 (2024).

4. Ostersehlt, L. M. et al. DNA-PAINT MINFLUX nanoscopy. Nat. Methods 1G, 1072–1075 (2022).

5. Shaib, A. H. et al. One-step nanoscale expansion microscopy reveals individual protein shapes. Nat. Biotechnol. 10.1038/s41587-024-02431-9 (2024)

6. Bond, C., Santiago-Ruiz, A. N., Tang, Q. C, Lakadamyali, M. Technological advances in super-resolution microscopy to study cellular processes. Mol. Cell 82, 315–332 (2022).

7. Masullo, L. A., Szalai, A. M., Lopez, L. F. C, Stefani, F. D. Fluorescence nanoscopy at the sub-10 nm scale. Biophys. Rev. 13, 1101–1112 (2021).

8. Helmerich, D. A. et al. Photoswitching fingerprint analysis bypasses the 10-nm resolution barrier. Nat. Methods **1G**, 986–994 (2022).

9. Ebert, V., Sauer, M. C, Doose, S. Energy transfer leaves fingerprints in cyanine photoswitching behavior. PLOS Comput. Biol. 22, e1014322 (2026).

10. Früh, S. M., et al. Site-Specifically-Labeled Antibodies for Super-Resolution Microscopy Reveal In Situ Linkage Errors. ACS Nano 15, 12161–12170 (2021).

11. Helmerich, D. A. et al. Impact of Docking Strand Design on Spatial Resolution in DNA-Points Accumulation for Imaging in Nanoscale Topography. ChemPhysChem 27, e202500803 (2026).

12. Zwettler, F. U. et al. Molecular resolution imaging by post-labeling expansion single-molecule localization microscopy (Ex-SMLM). Nat. Commun. 11, 3388 (2020).

13. Thevathasan, J. V. et al. Nuclear pores as versatile reference standards for quantitative superresolution microscopy. Nat. Methods 16, 1045–1053 (2019).

14. Scheckenbach, M., Bauer, J., Zähringer, J., Selbach, F. C, Tinnefeld, P. DNA origami nanorulers and emerging reference structures. APL Mater. 8, 110902 (2020).

15. Akbari, E. et al. Engineering Cell Surface Function with DNA Origami. Adv. Mater. **2G**, 1703632 (2017).

16. Mills, A., Aissaoui, N., Finkel, J., Elezgaray, J. C, Bellot, G. Mechanical DNA Origami to Investigate Biological Systems. *Adv*. Biol. 7, 2200224 (2023).

17. Helmerich, D. A. et al. PCNA as Protein-Based Nanoruler for Sub-10 nm Fluorescence Imaging. Adv. Mater. 2310104 (2023) doi:10.1002/adma.202310104.

18. Smith, E. R. et al. A Universal Protein Ladder for Standardization of Diverse FRET Assays. Adv. Sci. **n/a**, e76672.

19. Correnti, C. E. et al. Engineering and functionalization of large circular tandem repeat protein nanoparticles. Nat. Struct. Mol. Biol. 27, 342–350 (2020).

20. Eddins, A. J. et al. Truncation-Free Genetic Code Expansion with Tetrazine Amino Acids for Quantitative Protein Ligations. Bioconjug. Chem. 34, 2243–2254 (2023).

21. Xu, H. et al. Re-exploration of the Codon Context Effect on Amber Codon-Guided Incorporation of Noncanonical Amino Acids in Escherichia coli by the Blue–White Screening Assay. ChemBioChem 17, 1250–1256 (2016).

22. Lajoie, M. J. et al. Genomically Recoded Organisms Expand Biological Functions. Science 342, 357–360 (2013).

23. Bartoschek, M. D. et al. Identification of permissive amber suppression sites for efficient non-canonical amino acid incorporation in mammalian cells. Nucleic Acids Res. 4G, e62 (2021).

24. Damstra, H. G. et al. Visualizing cellular and tissue ultrastructure using Ten-fold Robust Expansion Microscopy (TREx). eLife 11, e73775 (2022).

25. Amiram, M. et al. Evolution of translation machinery in recoded bacteria enables multi-site incorporation of nonstandard amino acids. Nat. Biotechnol. 33, 1272–1279 (2015).

26. Streit, M. et al. Optimized genetic code expansion technology for time-dependent induction of adhesion GPCR-ligand engagement. Protein Sci. 32, e4614 (2023).

27. Li, D., Liu, L. C Li, W.-H. Genetic Targeting of a Small Fluorescent Zinc Indicator to Cell Surface for Monitoring Zinc Secretion. ACS Chem. Biol. 10, 1054–1063 (2015).

28. Wolter, S. rapidSTORM: accurate, fast open-source software for localization microscopy. Nat Methods **G**, (2012).

29. Ovesný, M., Køížek, P., Borkovec, J., Švindrych, Z. C, Hagen, G. M. ThunderSTORM: a comprehensive ImageJ plug-in for PALM and STORM data analysis and super-resolution imaging. Bioinformatics 30, 2389–2390 (2014).

