## Supplementary material for "Programmable Protein Reference Standards for benchmarking sub-10 nm Fluorescence Microscopy": cTRP_PicoRulers_Supplementary_Information_v1

**Table S1.** Characteristics of different variants of recombinant wild-type cTRPs

| cTRP variants | Amino acid sequences* | $\varepsilon_{280}$ ($M^{-1}$ $cm^{-1}$) | MW (g/mol) |
|---|---|---|---|
| **1** | MASSHHHHHHSSGLVPRGSSMGLDPEAIKAAAELGKAGISSEEILELLRAAHEL<br>GLDPEAIKAAAELGKAGISSEEILELLRAAHELGLDPEAIKAAAELGKAGISSEEILELLRAAHEL<br>GLDPEAIKAAAELGKAGISSEEILELLRAAHELGLDPEAIKAAAELGKAGISSEEILELLRAAHEL<br>GLDPEAIKAAAELGKAGISSEEILELLRAAHELGLDPEAIKAAAELGKAGISSEEILELLRAAHEL<br>GLDPEAIKAAAELGKAGISSEEILELLRAAHELGLDPEAIKAAAELGKAGISSEEILELLRAAHEL<br>GLDPEAIKAAAELGKAGISSEEILELLRAAHELGLDPEAIKAAAELGKAGISSEEILELLRAAHEL<br>GLDPEAIKAAAELGKAGISSEEILELLRAAHELGLDPEAIKAAAELGKAGISSEEILELLRAAHEL<br>GLDPEAIKAAAELGKAGISSEEILELLRAAHELGLDPEAIKAAAELGKAGISSEEILELLRAAHEL<br>GLDPEAIKAAAELGKAGISSEEILELLRAAHELGLDPEAIKAAAELGKAGISSEEILELLRAAHEL<br>GLDPEAIKAAAELGKAGISSEEILELLRAAHELGLDPEAIKAAAELGKAGISSEEILELLRAAHEL<br>GLDPEAIKAAAELGKAGISSEEILELLRAAHELGLDPEAIKAAAELGKAGISSEEILELLRAAHEL<br>GLDPEAIKAAAELGKAGISSEEILELLRAAHELGLDPEAIKAAAELGKAGISSEEILELLRAAHEL<br>GLDPEAIKAAAELGKAGISSEEILELLRAAHELGGW | 5500 | 84125.77 |
| **2** | MASSHHHHHHSSGLVPRGSSMGLDCEAIKAAAELGKAGISSEEILELLRAAHEL<br>GLDPEAIKAAAELGKAGISSEEILELLRAAHELGLDPEAIKAAAELGKAGISSEEILELLRAAHEL<br>GLDPEAIKAAAELGKAGISSEEILELLRAAHELGLDPEAIKAAAELGKAGISSEEILELLRAAHEL<br>GLDPEAIKAAAELGKAGISSEEILELLRAAHELGLDPEAIKAAAELGKAGISSEEILELLRAAHEL<br>GLDPEAIKAAAELGKAGISSEEILELLRAAHELGLDPEAIKAAAELGKAGISSEEILELLRAAHEL<br>GLDPEAIKAAAELGKAGISSEEILELLRAAHELGLDPEAIKAAAELGKAGISSEEILELLRAAHEL<br>GLDPECIKAAAELGKAGISSEEILELLRAAHELGGW | 11250[a]<br>5500[b] | 86774.74[a]<br>43389.37[b] |
| **3** | MASSHHHHHHSSGLVPRGSSMGLDCEAIKAAAELGKAGISSEEILELLRAAHEL<br>GLDPEAIKAAAELGKAGISSEEILELLRAAHELGLDPEAIKAAAELGKAGISSEEILELLRAAHVL<br>GLDPEAIKAAAELGKAGISSEEILELLVAAHLLGLDPEAIKAAAELGKAGISSEEILELLRAAHEL<br>GLDPEAIKAAAELGKAGISSEEILELLRAAHELGLDPEAIKAAAELGKAGISSEEILELLRAAHEL<br>GLDPECIKAAAELGKAGISSIEILELLRAAHELGGW | 16875[a]<br>5500[b] | 89030.79[a]<br>29678.93[b] |
| **4** | MASSHHHHHHSSGLVPRGSSMGLDCEAIKAAAELGKAGISSEEILELLRAAHEL<br>GLDPEAIKAAAELGKAGISSEEILELLRAAHELGLDPEAIKAAAELGKAGISSEEILELLRAAHEL<br>GLDPEAIKAAAELGKAGISSEEILELLRAAHELGLDPEAIKAAAELGKAGISSEEILELLRAAHEL<br>GLDPECIKAAAELGKAGISSEEILELLRAAHELGGW* | 22500[a]<br>5500[b] | 92000.52[a]<br>23002.13[b] |

His6X tag (green), thrombin cleavage-site (cyan), and cysteine residues responsible for disulfide-linked ring closure (yellow) are highlighted with different colors. Molecular weights and extinction coefficients at 280 nm are indicated as follows: [a] for the fully assembled ring, and [b] for a single subunit (monomer).

**Table S2.** Characteristics of different variants of recombinant clickable cTRPs

| cTRP variants | Amino acid sequences* | $\varepsilon_{280}$ ($M^{-1}$ $cm^{-1}$) | MW (g/mol) |
|---|---|---|---|
| **cTRP-1** | MASSHHHHHHSSGLVPRGSSMGLDCEAIKAAAELGKAGISSEEILELLRAAHEL<br>GLDPEAIKAAAELGKAGISSEEILELLRAAHELGLDPEAIKAAAELGKAGISSEEILELLRAAHEL<br>GLDPEAIKAAAELGKAGISSEEILELLRAAHELGLDPEAIKAAAELGKAGISSEEILELLRAAHEL<br>GLDPEAIKAAAELGKAGISSEEILELLRAAHELGLDPEAIKAAAELGKAGISSEEILELLRAAHEL<br>GLDPEAIKAAAELGKAGISSEEILELLRAAHELGLDPEAIKAAAELGKAGISSEEILELLRAAHEL<br>GLDPEAIKAAAELGKAGISSEEILELLRAAHELGLDPEAIKAAAELGKAGISSEEILELLRAAHEL<br>GLDPEAIKAAAELGKAGISSEEILELLRAAHELGSNXTGLDPEAIKAAAELGKAGISSEEILELLRAAHEL<br>GLDPEAIKAAAELGKAGISSEEILELLRAAHELGLDPEAIKAAAELGKAGISSEEILELLRAAHEL<br>GLDPEAIKAAAELGKAGISSEEILELLRAAHELGLDPEAIKAAAELGKAGISSEEILELLRAAHEL<br>GLDPEAIKAAAELGKAGISSEEILELLRAAHELGLDPEAIKAAAELGKAGISSEEILELLRAAHEL<br>GLDPEAIKAAAELGKAGISSEEILELLRAAHELGLDPEAIKAAAELGKAGISSEEILELLRAAHEL<br>GLDPEAIKAAAELGKAGISSEEILELLRAAHELGLDPEAIKAAAELGKAGISSEEILELLRAAHEL<br>GLDPECIKAAAELGKAGISSEEILELLRAAHELGGWGGSGGLEVLFQGPWSHPQFEK | 24960 | 86835.73 |
| **cTRP-2** | MASSHHHHHHSSGLVPRGSSMGLDCEAIKAAAELGKAGISSEEILELLRAAHEL<br>GLDPEAIKAAAELGKAGISSEEILELLRAAHELGLDPEAIKAAAELGKAGISSEEILELLRAAHEL<br>GLDPEAIKAAAELGKAGISSEEILELLRAAHELGLDPEAIKAAAELGKAGISSEEILELLRAAHEL<br>GLDPEAIKAAAELGKAGISSEEILELLRAAHELGLDPEAIKAAAELGKAGISSEEILELLRAAHELGSNXT<br>GLDPEAIKAAAELGKAGISSEEILELLRAAHELGLDPEAIKAAAELGKAGISSEEILELLRAAHEL<br>GLDPEAIKAAAELGKAGISSEEILELLRAAHELGLDPEAIKAAAELGKAGISSEEILELLRAAHEL<br>GLDPEAIKAAAELGKAGISSEEILELLRAAHELGLDPEAIKAAAELGKAGISSEEILELLRAAHEL<br>GLDPEAIKAAAELGKAGISSEEILELLRAAHELGLDPEAIKAAAELGKAGISSEEILELLRAAHEL<br>GLDPEAIKAAAELGKAGISSEEILELLRAAHELGLDPEAIKAAAELGKAGISSEEILELLRAAHEL<br>GLDPEAIKAAAELGKAGISSEEILELLRAAHELGLDPEAIKAAAELGKAGISSEEILELLRAAHELGSNXT<br>GLDPEAIKAAAELGKAGISSEEILELLRAAHELGLDPEAIKAAAELGKAGISSEEILELLRAAHEL<br>GLDPEAIKAAAELGKAGISSEEILELLRAAHELGLDPEAIKAAAELGKAGISSEEILELLRAAHEL<br>GLDPECIKAAAELGKAGISSEEILELLRAAHELGGWGGSGGLEVLFQGPWSHPQFEK | 38645 | 87632.56 |
| **cTRP-3** | MASSHHHHHHSSGLVPRGSSMGLDCEAIKAAAELGKAGISSEEILELLRAAHEL<br>GLDPEAIKAAAELGKAGISSEEILELLRAAHELGLDPEAIKAAAELGKAGISSEEILELLRAAHEL<br>GLDPEAIKAAAELGKAGISSEEILELLRAAHELGSNXTGLDPEAIKAAAELGKAGISSEEILELLRAAHEL<br>GLDPEAIKAAAELGKAGISSEEILELLRAAHELGLDPEAIKAAAELGKAGISSEEILELLRAAHEL<br>GLDPEAIKAAAELGKAGISSEEILELLRAAHELGLDPEAIKAAAELGKAGISSEEILELLRAAHEL<br>GLDPEAIKAAAELGKAGISSEEILELLRAAHELGLDPEAIKAAAELGKAGISSEEILELLRAAHEL<br>GLDPEAIKAAAELGKAGISSEEILELLRAAHELGSNXTGLDPEAIKAAAELGKAGISSEEILELLRAAHEL<br>GLDPEAIKAAAELGKAGISSEEILELLRAAHELGLDPEAIKAAAELGKAGISSEEILELLRAAHEL<br>GLDPEAIKAAAELGKAGISSEEILELLRAAHELGLDPEAIKAAAELGKAGISSEEILELLRAAHEL<br>GLDPEAIKAAAELGKAGISSEEILELLRAAHELGLDPEAIKAAAELGKAGISSEEILELLRAAHEL<br>GLDPEAIKAAAELGKAGISSEEILELLRAAHELGSNXTGLDPEAIKAAAELGKAGISSEEILELLRAAHEL<br>GLDPEAIKAAAELGKAGISSEEILELLRAAHELGLDPEAIKAAAELGKAGISSEEILELLRAAHEL<br>GLDPECIKAAAELGKAGISSEEILELLRAAHELGGWGGSGGLEVLFQGPWSHPQFEK | 52205 | 88247.17 |
| **cTRP-4** | MASSHHHHHHSSGLVPRGSSMGLDCEAIKAAAELGKAGISSEEILELLRAAHEL<br>GLDPEAIKAAAELGKAGISSEEILELLRAAHELGLDPEAIKAAAELGKAGISSEEILELLRAAHELGSNXT<br>GLDPEAIKAAAELGKAGISSEEILELLRAAHELGLDPEAIKAAAELGKAGISSEEILELLRAAHEL<br>GLDPEAIKAAAELGKAGISSEEILELLRAAHELGLDPEAIKAAAELGKAGISSEEILELLRAAHEL<br>GLDPEAIKAAAELGKAGISSEEILELLRAAHELGLDPEAIKAAAELGKAGISSEEILELLRAAHELGSNXT<br>GLDPEAIKAAAELGKAGISSEEILELLRAAHELGLDPEAIKAAAELGKAGISSEEILELLRAAHEL<br>GLDPEAIKAAAELGKAGISSEEILELLRAAHELGLDPEAIKAAAELGKAGISSEEILELLRAAHEL<br>GLDPEAIKAAAELGKAGISSEEILELLRAAHELGLDPEAIKAAAELGKAGISSEEILELLRAAHEL<br>GSNXTGLDPEAIKAAAELGKAGISSEEILELLRAAHELGLDPEAIKAAAELGKAGISSEEILELLRAAHEL<br>GLDPEAIKAAAELGKAGISSEEILELLRAAHELGLDPEAIKAAAELGKAGISSEEILELLRAAHEL<br>GLDPEAIKAAAELGKAGISSEEILELLRAAHELGLDPEAIKAAAELGKAGISSEEILELLRAAHELGSNXT<br>GLDPEAIKAAAELGKAGISSEEILELLRAAHELGLDPEAIKAAAELGKAGISSEEILELLRAAHEL<br>GLDPECIKAAAELGKAGISSEEILELLRAAHELGGWGGSGGLEVLFQGPWSHPQFEK | 65765 | 88861.78 |

| | | | |
|---|---|---|---|
| **cTRP-6** | MASSHHHHHHSSGLVPRGSSMGLDCEAIKAAAELGKAGISSEEILELLRAAHEL<br>GLDPEAIKAAAELGKAGISSEEILELLRAAHELGSNXTGLDPEAIKAAAELGKAGISSEEILELLRAAHEL<br>GLDPEAIKAAAELGKAGISSEEILELLRAAHELGLDPEAIKAAAELGKAGISSEEILELLRAAHEL<br>GLDPEAIKAAAELGKAGISSEEILELLRAAHELGSNXTGLDPEAIKAAAELGKAGISSEEILELLRAAHEL<br>GLDPEAIKAAAELGKAGISSEEILELLRAAHELGLDPEAIKAAAELGKAGISSEEILELLRAAHEL<br>GLDPEAIKAAAELGKAGISSEEILELLRAAHELGSNXTGLDPEAIKAAAELGKAGISSEEILELLRAAHEL<br>GLDPEAIKAAAELGKAGISSEEILELLRAAHELGLDPEAIKAAAELGKAGISSEEILELLRAAHEL<br>GLDPEAIKAAAELGKAGISSEEILELLRAAHELGSNXTGLDPEAIKAAAELGKAGISSEEILELLRAAHEL<br>GLDPEAIKAAAELGKAGISSEEILELLRAAHELGLDPEAIKAAAELGKAGISSEEILELLRAAHEL<br>GLDPEAIKAAAELGKAGISSEEILELLRAAHELGSNXTGLDPEAIKAAAELGKAGISSEEILELLRAAHEL<br>GLDPEAIKAAAELGKAGISSEEILELLRAAHELGLDPEAIKAAAELGKAGISSEEILELLRAAHEL<br>GLDPEAIKAAAELGKAGISSEEILELLRAAHELGSNXTGLDPEAIKAAAELGKAGISSEEILELLRAAHEL<br>GLDPECIKAAAELGKAGISSEEILELLRAAHELGGWGGSGGLEVLFQGPWSHPQFEK | 92885 | 90091.01 |

*All cTRP PicoRulers were expressed with clickable ncAA ethyl tetrazine (X = Tet2Et or pAzF). Functional elements are color-coded as follows: His6X tag (green), thrombin cleavage site (cyan), disulfide-forming cysteine residues (yellow), ethyl tetrazine (teal), HRV-3C protease cleavage site (gray), and Strep-tag II (magenta).

**Table S3.** Amino acid sequences of genetically encoded clickable membrane-displayed cTRPs

| cTRP variants | Amino acid sequences* |
|---|---|
| **cTRP-1** | MRAWIFFLLCLAGRALAAPLAGTHHHHHHGSGDYKDDDDKGSGLVPRGSSM<br>GLDCEAIKAAAELGKAGISSEEILELLRAAHELGLDPEAIKAAAELGKAGISSEEILELLRAAHEL<br>GLDPEAIKAAAELGKAGISSEEILELLRAAHELGLDPEAIKAAAELGKAGISSEEILELLRAAHEL<br>GLDPEAIKAAAELGKAGISSEEILELLRAAHELGLDPEAIKAAAELGKAGISSEEILELLRAAHEL<br>GLDPEAIKAAAELGKAGISSEEILELLRAAHELGLDPEAIKAAAELGKAGISSEEILELLRAAHEL<br>GLDPEAIKAAAELGKAGISSEEILELLRAAHELGLDPEAIKAAAELGKAGISSEEILELLRAAHEL<br>GLDPEAIKAAAELGKAGISSEEILELLRAAHELGLDPEAIKAAAELGKAGISSEEILELLRAAHELGSGXL<br>GLDPEAIKAAAELGKAGISSEEILELLRAAHELGLDPEAIKAAAELGKAGISSEEILELLRAAHEL<br>GLDPEAIKAAAELGKAGISSEEILELLRAAHELGLDPEAIKAAAELGKAGISSEEILELLRAAHEL<br>GLDPEAIKAAAELGKAGISSEEILELLRAAHELGLDPEAIKAAAELGKAGISSEEILELLRAAHEL<br>GLDPEAIKAAAELGKAGISSEEILELLRAAHELGLDPEAIKAAAELGKAGISSEEILELLRAAHEL<br>GLDPEAIKAAAELGKAGISSEEILELLRAAHELGLDPEAIKAAAELGKAGISSEEILELLRAAHEL<br>GLDPEAIKAAAELGKAGISSEEILELLRAAHELGLDPECIKAAAELGKAGISSEEILELLRAAHEL<br>GGWTGSEAAAKEAAAKEAAAKEAAAKEAAAKEAAAKGSLEVLFQGPGGGGSGGGGSGGGGS<br>QLENGGISLLVQNTSWMLLLLLSLSLLQALDFISL |
| **cTRP-2** | MRAWIFFLLCLAGRALAAPLAGTHHHHHHGSGDYKDDDDKGSGLVPRGSSM<br>GLDCEAIKAAAELGKAGISSEEILELLRAAHELGLDPEAIKAAAELGKAGISSEEILELLRAAHEL<br>GLDPEAIKAAAELGKAGISSEEILELLRAAHELGLDPEAIKAAAELGKAGISSEEILELLRAAHEL<br>GLDPEAIKAAAELGKAGISSEEILELLRAAHELGLDPEAIKAAAELGKAGISSEEILELLRAAHEL<br>GLDPEAIKAAAELGKAGISSEEILELLRAAHELGSGXLGLDPEAIKAAAELGKAGISSEEILELLRAAHEL<br>GLDPEAIKAAAELGKAGISSEEILELLRAAHELGLDPEAIKAAAELGKAGISSEEILELLRAAHEL<br>GLDPEAIKAAAELGKAGISSEEILELLRAAHELGLDPEAIKAAAELGKAGISSEEILELLRAAHEL<br>GLDPEAIKAAAELGKAGISSEEILELLRAAHELGLDPEAIKAAAELGKAGISSEEILELLRAAHEL<br>GLDPEAIKAAAELGKAGISSEEILELLRAAHELGLDPEAIKAAAELGKAGISSEEILELLRAAHEL<br>GLDPEAIKAAAELGKAGISSEEILELLRAAHELGLDPEAIKAAAELGKAGISSEEILELLRAAHEL<br>GLDPEAIKAAAELGKAGISSEEILELLRAAHELGSGXLGLDPEAIKAAAELGKAGISSEEILELLRAAHEL<br>GLDPEAIKAAAELGKAGISSEEILELLRAAHELGLDPEAIKAAAELGKAGISSEEILELLRAAHEL<br>GLDPEAIKAAAELGKAGISSEEILELLRAAHELGLDPECIKAAAELGKAGISSEEILELLRAAHEL<br>GGWTGSEAAAKEAAAKEAAAKEAAAKEAAAKEAAAKGSLEVLFQGPGGGGSGGGGSGGGGS<br>QLENGGISLLVQNTSWMLLLLLSLSLLQALDFISL |
| **cTRP-2B** | MRAWIFFLLCLAGRALAAPLAGTHHHHHHGSGDYKDDDDKGSGLVPRGSSM<br>GLDCEAIKAAAELGKAGISSEEILELLRAAHELGLDPEAIKAAAELGKAGISSEEILELLRAAHEL<br>GLDPEAIKAAAELGKAGISSEEILELLRAAHELGLDPEAIKAAAELGKAGISSEEILELLRAAHEL<br>GLDPEAIKAAAELGKAGISSEEILELLRAAHELGLDPEAIKAAAELGKAGISSEEILELLRAAHEL<br>GLDPEAIKAAAELGKAGISSEEILELLRAAHELGLDPEAIKAAAELGKAGISSEEILELLRAAHEL<br>GLDPEAIKAAAELGKAGISSEEILELLRAAHELGLDPEAIKAAAELGKAGISSEEILELLRAAHEL<br>GLDPEAIKAAAELGKAGISSEEILELLRAAHELGLDPEAIKAAAELGKAGISSEEILELLRAAHELGSGXL<br>GLDPEAIKAAAELGKAGISSEEILELLRAAHELGLDPEAIKAAAELGKAGISSEEILELLRAAHEL<br>GLDPEAIKAAAELGKAGISSEEILELLRAAHELGSGXLGLDPEAIKAAAELGKAGISSEEILELLRAAHEL<br>GLDPEAIKAAAELGKAGISSEEILELLRAAHELGLDPEAIKAAAELGKAGISSEEILELLRAAHEL<br>GLDPEAIKAAAELGKAGISSEEILELLRAAHELGLDPEAIKAAAELGKAGISSEEILELLRAAHEL<br>GLDPEAIKAAAELGKAGISSEEILELLRAAHELGLDPEAIKAAAELGKAGISSEEILELLRAAHEL<br>GLDPEAIKAAAELGKAGISSEEILELLRAAHELGLDPECIKAAAELGKAGISSEEILELLRAAHEL<br>GGWTGSEAAAKEAAAKEAAAKEAAAKEAAAKEAAAKGSLEVLFQGPGGGGSGGGGSGGGGS<br>QLENGGISLLVQNTSWMLLLLLSLSLLQALDFISL |
| **cTRP-3** | MRAWIFFLLCLAGRALAAPLAGTHHHHHHGSGDYKDDDDKGSGLVPRGSSM<br>GLDCEAIKAAAELGKAGISSEEILELLRAAHELGLDPEAIKAAAELGKAGISSEEILELLRAAHEL<br>GLDPEAIKAAAELGKAGISSEEILELLRAAHELGLDPEAIKAAAELGKAGISSEEILELLRAAHELGSGXL<br>GLDPEAIKAAAELGKAGISSEEILELLRAAHELGLDPEAIKAAAELGKAGISSEEILELLRAAHEL<br>GLDPEAIKAAAELGKAGISSEEILELLRAAHELGLDPEAIKAAAELGKAGISSEEILELLRAAHEL<br>GLDPEAIKAAAELGKAGISSEEILELLRAAHELGLDPEAIKAAAELGKAGISSEEILELLRAAHEL<br>GLDPEAIKAAAELGKAGISSEEILELLRAAHELGLDPEAIKAAAELGKAGISSEEILELLRAAHELGSGXL<br>GLDPEAIKAAAELGKAGISSEEILELLRAAHELGLDPEAIKAAAELGKAGISSEEILELLRAAHEL<br>GLDPEAIKAAAELGKAGISSEEILELLRAAHELGLDPEAIKAAAELGKAGISSEEILELLRAAHEL<br>GLDPEAIKAAAELGKAGISSEEILELLRAAHELGLDPEAIKAAAELGKAGISSEEILELLRAAHEL<br>GLDPEAIKAAAELGKAGISSEEILELLRAAHELGLDPEAIKAAAELGKAGISSEEILELLRAAHELGSGXL<br>GLDPEAIKAAAELGKAGISSEEILELLRAAHELGLDPEAIKAAAELGKAGISSEEILELLRAAHEL<br>GLDPEAIKAAAELGKAGISSEEILELLRAAHELGLDPECIKAAAELGKAGISSEEILELLRAAHEL<br>GGWTGSEAAAKEAAAKEAAAKEAAAKEAAAKEAAAKGSLEVLFQGPGGGGSGGGGSGGGGS<br>QLENGGISLLVQNTSWMLLLLLSLSLLQALDFISL |

| | |
|---|---|
| **cTRP-3B** | MRAWIFFLLCLAGRALAAPLAGTHHHHHHGSGDYKDDDDKGSGLVPRGSSM<br>GLDCEAIKAAAELGKAGISSEEILELLRAAHELGLDPEAIKAAAELGKAGISSEEILELLRAAHEL<br>GLDPEAIKAAAELGKAGISSEEILELLRAAHELGLDPEAIKAAAELGKAGISSEEILELLRAAHEL<br>GLDPEAIKAAAELGKAGISSEEILELLRAAHELGLDPEAIKAAAELGKAGISSEEILELLRAAHEL<br>GLDPEAIKAAAELGKAGISSEEILELLRAAHELGLDPEAIKAAAELGKAGISSEEILELLRAAHEL<br>GLDPEAIKAAAELGKAGISSEEILELLRAAHELGSGXLGLDPEAIKAAAELGKAGISSEEILELLRAAHEL<br>GLDPEAIKAAAELGKAGISSEEILELLRAAHELGLDPEAIKAAAELGKAGISSEEILELLRAAHELGSGXL<br>GLDPEAIKAAAELGKAGISSEEILELLRAAHELGLDPEAIKAAAELGKAGISSEEILELLRAAHEL<br>GLDPEAIKAAAELGKAGISSEEILELLRAAHELGSGXLGLDPEAIKAAAELGKAGISSEEILELLRAAHEL<br>GLDPEAIKAAAELGKAGISSEEILELLRAAHELGLDPEAIKAAAELGKAGISSEEILELLRAAHEL<br>GLDPEAIKAAAELGKAGISSEEILELLRAAHELGLDPEAIKAAAELGKAGISSEEILELLRAAHEL<br>GLDPEAIKAAAELGKAGISSEEILELLRAAHELGLDPEAIKAAAELGKAGISSEEILELLRAAHEL<br>GLDPEAIKAAAELGKAGISSEEILELLRAAHELGLDPECIKAAAELGKAGISSEEILELLRAAHEL<br>GGWTGSEAAAKEAAAKEAAAKEAAAKEAAAKEAAAKGSLEVLFQGPGGGGSGGGGSGGGGS<br>QLENGGISLLVQNTSWMLLLLLSLSLLQALDFISL |
| **cTRP-4** | MRAWIFFLLCLAGRALAAPLAGTHHHHHHGSGDYKDDDDKGSGLVPRGSSM<br>GLDCEAIKAAAELGKAGISSEEILELLRAAHELGLDPEAIKAAAELGKAGISSEEILELLRAAHEL<br>GLDPEAIKAAAELGKAGISSEEILELLRAAHELGSGXLGLDPEAIKAAAELGKAGISSEEILELLRAAHEL<br>GLDPEAIKAAAELGKAGISSEEILELLRAAHELGLDPEAIKAAAELGKAGISSEEILELLRAAHEL<br>GLDPEAIKAAAELGKAGISSEEILELLRAAHELGLDPEAIKAAAELGKAGISSEEILELLRAAHEL<br>GLDPEAIKAAAELGKAGISSEEILELLRAAHELGSGXLGLDPEAIKAAAELGKAGISSEEILELLRAAHEL<br>GLDPEAIKAAAELGKAGISSEEILELLRAAHELGLDPEAIKAAAELGKAGISSEEILELLRAAHEL<br>GLDPEAIKAAAELGKAGISSEEILELLRAAHELGLDPEAIKAAAELGKAGISSEEILELLRAAHEL<br>GLDPEAIKAAAELGKAGISSEEILELLRAAHELGSGXLGLDPEAIKAAAELGKAGISSEEILELLRAAHEL<br>GLDPEAIKAAAELGKAGISSEEILELLRAAHELGLDPEAIKAAAELGKAGISSEEILELLRAAHEL<br>GLDPEAIKAAAELGKAGISSEEILELLRAAHELGLDPEAIKAAAELGKAGISSEEILELLRAAHEL<br>GLDPEAIKAAAELGKAGISSEEILELLRAAHELGSGXLGLDPEAIKAAAELGKAGISSEEILELLRAAHEL<br>GLDPEAIKAAAELGKAGISSEEILELLRAAHELGLDPECIKAAAELGKAGISSEEILELLRAAHEL<br>GGWTGSEAAAKEAAAKEAAAKEAAAKEAAAKEAAAKGSLEVLFQGPGGGGSGGGGSGGGGS<br>QLENGGISLLVQNTSWMLLLLLSLSLLQALDFISL |
| **cTRP-6** | MRAWIFFLLCLAGRALAAPLAGTHHHHHHGSGDYKDDDDKGSGLVPRGSSM<br>GLDCEAIKAAAELGKAGISSEEILELLRAAHELGLDPEAIKAAAELGKAGISSEEILELLRAAHELGSGXL<br>GLDPEAIKAAAELGKAGISSEEILELLRAAHELGLDPEAIKAAAELGKAGISSEEILELLRAAHEL<br>GLDPEAIKAAAELGKAGISSEEILELLRAAHELGLDPEAIKAAAELGKAGISSEEILELLRAAHELGSGXL<br>GLDPEAIKAAAELGKAGISSEEILELLRAAHELGLDPEAIKAAAELGKAGISSEEILELLRAAHEL<br>GLDPEAIKAAAELGKAGISSEEILELLRAAHELGLDPEAIKAAAELGKAGISSEEILELLRAAHELGSGXL<br>GLDPEAIKAAAELGKAGISSEEILELLRAAHELGLDPEAIKAAAELGKAGISSEEILELLRAAHEL<br>GLDPEAIKAAAELGKAGISSEEILELLRAAHELGLDPEAIKAAAELGKAGISSEEILELLRAAHELGSGXL<br>GLDPEAIKAAAELGKAGISSEEILELLRAAHELGLDPEAIKAAAELGKAGISSEEILELLRAAHEL<br>GLDPEAIKAAAELGKAGISSEEILELLRAAHELGLDPEAIKAAAELGKAGISSEEILELLRAAHELGSGXL<br>GLDPEAIKAAAELGKAGISSEEILELLRAAHELGLDPEAIKAAAELGKAGISSEEILELLRAAHEL<br>GLDPEAIKAAAELGKAGISSEEILELLRAAHELGLDPEAIKAAAELGKAGISSEEILELLRAAHELGSGXL<br>GLDPEAIKAAAELGKAGISSEEILELLRAAHELGLDPECIKAAAELGKAGISSEEILELLRAAHEL<br>GGWTGSEAAAKEAAAKEAAAKEAAAKEAAAKEAAAKGSLEVLFQGPGGGGSGGGGSGGGGS<br>QLENGGISLLVQNTSWMLLLLLSLSLLQALDFISL |

*All cTRP PicoRulers were expressed with clickable ncAA trans-cyclooctene (X = TCO*A). Functional elements are color-coded as follows: Signaling peptide (olive green), His6X tag (green), FLAG tag (magenta), thrombin cleavage site (cyan), disulfide-forming cysteine residues (yellow), trans-cyclooctene (teal), rigid helix linker $(EAAAK)_6$ (dark green), HRV-3C protease cleavage site (gray), flexible linker $(G_4S)_3$ (red) and GPI (dark blue).
